# The Implant-Induced Foreign Body Response is Limited by CD13-Dependent Regulation of Ubiquitination of Fusogenic Proteins

**DOI:** 10.1101/2023.09.20.558631

**Authors:** Mallika Ghosh, Fraser McGurk, Rachael Norris, Andy Dong, Sreenidhi Nair, Evan Jellison, Patrick Murphy, RajKumar Verma, Linda H Shapiro

## Abstract

Implanted medical devices from artificial heart valves, arthroscopic joints to implantable sensors often induce a Foreign Body Response (FBR), a form of chronic inflammation resulting from the inflammatory reaction to a persistent foreign stimulus. The FBR is characterized by a subset of multinucleated giant cells (MGCs) formed by macrophage fusion, the Foreign Body Giant cells (FBGCs), accompanied by inflammatory cytokines, matrix deposition and eventually, deleterious fibrotic implant encapsulation. Despite efforts to improve biocompatibility, implant-induced FBR persists, compromising the utility of devices and making efforts to control the FBR imperative for long-term function. Controlling macrophage fusion in FBGC formation presents a logical target to prevent implant failure, but the actual contribution of FBGCs to FBR-induced damage is controversial. CD13 is a molecular scaffold and *in vitro* induction of CD13^KO^ bone-marrow progenitors generates many more MGCs than WT, suggesting CD13 regulates macrophage fusion. Moreover, in the mesh implant model of FBR, CD13^KO^ mice produced significantly more peri-implant FBGCs with enhanced TGFβ expression and increased collagen deposition vs. WT. Pre-fusion, increased protrusion and microprojection formation accompanies hyperfusion in the absence of CD13. Expression of fusogenic proteins driving cell-cell fusion was aberrantly sustained at high levels in CD13^KO^ MGCs, which we show is due to a novel CD13 function, regulating ubiquitin/proteasomal protein degradation. By controlling protein degradation, CD13 becomes a physiologic brake preventing aberrant macrophage fusion and may be a novel therapeutic target to improve success of implanted medical devices. Furthermore, our data directly implicates FBGCs in the detrimental fibrosis that characterizes the FBR.

## Introduction

In mammals, both homotypic and heterotypic cell-cell fusion are crucial to multiple physiological and pathological processes (1). The mechanisms and proteins controlling cell-cell fusion in different cell types are remarkably diverse across the various systems in their structure, function and utilization of auxiliary pathways. However, some of the molecules responsible for merging individual membranes appear to be shared and thus participate in fusion of more than one cell type. Several of these key shared proteins have been identified, but our mechanistic understanding of precisely how these may promote cell-cell fusion is still largely theoretical and somewhat controversial, suggesting that additional proteins that drive fusion are yet to be discovered (2, 3). Clearly, identifying novel membrane-active fusion proteins and dissecting their underlying mechanisms will provide the clues to inform and direct further investigations into shared and unique mechanisms of membrane remodeling, membrane merger and cell-cell fusion.

Multinucleated giant cells (MGCs) are specialized cells of the immune system that result from the fusion of mononuclear myeloid cells to form a single, large, multinucleated cell with a shared cytoplasm. MGC formation is typified by osteoclast fusion during bone remodeling, but the immune response to therapeutic implantation of foreign materials such as medical devices, mesh supports, arthroscopic implants and biosensors also provokes a robust MGC response, the Foreign Body Response (FBR) (4–7). Initially, circulating proteins adsorb to the surface of the implant eliciting an acute and chronic inflammatory response mediated by cytokine release by infiltrating immune cells (6). Macrophages at the implant site fuse to form a type of specialized MGC called foreign body giant cells (FBGCCGs), the hallmark cell of the FBR. This is followed by fibrin matrix deposition, granulation tissue and fibrous capsule formation around the implanted biomaterial (6) leading to loss of host-implant communication and implant failure. While FBGCs have been shown to be consistently present at the site of implant, the relative contribution of FGBC in FBR and more specifically in fibrosis is poorly understood (8, 9). However, numerous investigations have linked FBGCs to the degradation of foreign particles through secretion of ROS, MMPs, and acids (10), and are thought to be a major source of cytokines and chemokines (11, 12), ROS (13, 14) and angiogenic mediators. To date, investigations to improve implant failure primarily have focused on improving the biocompatibility of medical devices or preventing the encapsulation of biomaterial, have been unsuccessful.

FBGC formation is a multi-step process that initially requires differentiation of precursors into macrophages which fuse to form active MGCs in response to cytokines IL-4+IL-13. Membrane fusion involves various membrane-associated processes such as membrane assembly (15–17), clustering of membrane molecules (18), endocytosis (19), adhesion (20, 21), cytoskeletal protrusion and rearrangement (22), leading to the characteristic multinucleated giant cell. Independent investigations have implicated specific molecules as fusion promoters, or fusogens, such as DC-STAMP (23), OC-STAMP (24), dynamin (19), tetraspanins (16, 25–27), ATP6v0d2 (28), syncytin-1 (Syn-1 (29), annexins (30) and S100 (31) proteins. However, how and if these proteins are part of a larger network has not been elucidated (3, 32), thus warranting further investigation to integrate these into a unifying mechanism.

CD13, is a multifunctional transmembrane aminopeptidase that we and others have shown to be involved in cell migration, actin cytoskeletal organization, cell-cell and cell-ECM adhesion, receptor mediated endocytosis and recycling (33–38). Recently, we have demonstrated a novel role of CD13 in osteoclastogenesis at the level of cell-cell fusion (39) where CD13^KO^ mice exhibited a low bone mass phenotype with increased osteoclast numbers per bone surface area, consistent with enhanced giant cell-mediated osteolysis and bone destruction. In the current study, we extend these observations and demonstrate that *in vitro* induction of fusion in CD13-deficient myeloid progenitors generated from bone marrow or thioglycolate-elicited peritoneal macrophages results in hyperfusion to generate MGCs that are considerably larger in size and contain many more nuclei than those from their wild type counterparts, suggesting that CD13 also regulates MGC formation at the stage of fusion. In an established murine model of FBR, the subcutaneous implantation of polyethylene mesh in CD13^KO^ mice produces significantly more FBGCs over time with a concomitant increase in levels of the serum cytokines IL-1β and TNF-α. Furthermore, collagen deposition and intracellular TGFβ expression levels in FBGCs were significantly enhanced in the peri-implant region in the CD13^KO^ compared to WT mice, suggesting that FBGCs directly contribute to the FBR. Mechanistically, the expression levels of the key fusion proteins, DCSTAMP, the tetraspanins CD9 and CD81 were sustained at high levels in CD13^KO^ compared to WT MGCs post-fusion by a ubiquitin/proteasomal degradation pathway with loss of association of fusogens with the scaffold protein, Cullin-4A, a component of the CUL4-RING E3 ubiquitin ligase (CRL4 E3 ligase) complex involved in protein turnover (40, 41). In addition, actin rearrangements are critical for the formation and stability of the fusion complex along with the several proteins including syncytins that are involved in the merging of the cells. Indeed, TEM analysis of myeloid cells undergoing fusion indicated that CD13^KO^ macrophages clearly produced more actin protrusions and microprojections compared to WT prior to fusion as early as d2 post fusion. Taken together, our study provides three novel insights into the mechanism of FBGC formation and the evolution of the FBR: 1) FBGCs do not only identify the FBR but play an active role in promoting FBR-dependent fibrosis; 2) CD13 regulates the cytoskeletal changes necessary for protrusion-dependent FBGC fusion; and 3) CD13 controls protein turnover by promoting the proper assembly of ubiquitination complexes. Taken together, CD13 acts as a brake to regulate physiologic macrophage fusion in two distinct lineages, osteoclasts and FBGCs, and may be a target for therapeutic intervention at early stages to control pathologic consequences of aberrant cell-cell fusion.

## Results

### CD13 Negatively Regulates Myeloid Cell Fusion

Previous studies from our laboratory and other investigations have established that CD13 is an essential mechanistic component of cellular functions required for cell-cell fusion (33, 38). More recently, we have demonstrated that compared to WT animals, CD13^KO^ mice had markedly reduced bone density with increased numbers of osteoclasts per bone surface (39). In the published work, mice of both genotypes showed equivalent osteoclast progenitor populations, bone formation and mineral apposition rates, suggesting a functional defect in CD13^KO^ osteoclasts. Indeed, cytokine-mediated osteoclastogenesis *in vitro* in CD13^KO^ myeloid progenitors generated larger osteoclasts with considerably more nuclei and significantly higher bone resorptive function than WT, indicating that CD13 controls osteoclastogenesis at the fusion stage. Extending these observations, we investigated whether CD13 also impacts fusion of a distinct lineage of fusogenic myeloid cells that derive from different progenitors, the multinucleated giant cells (42). We differentiated flow-sorted WT and CD13^KO^ BM-derived progenitors (CD3^-^,B220^-^ NK1.1^-^,CD11b^lo/-^,CD115^hi^,Ly6G^+^) into macrophages in the presence of M-CSF for 3d, followed by incubation in IL-4+IL-13 for an additional 3-5d to promote MGC differentiation and fusion (**Fig. 1A,D, S1A)**. Giemsa-stained cells in areas with comparable cell densities were analyzed in a blinded manner and the extent of MGC fusion at d3 post-induction represented as the fusion index [(total number of nuclei in fused cells per field /total number of nucleifield) × 100]. We found significant increases (2 fold) in MGC formation in CD13^KO^ cells compared to WT as early as d3 **(Fig. 1D)**, suggesting that the presence of CD13 limits or restricts the fusion process. However, the total number of nuclei/dish was equivalent between genotypes, confirming that this accelerated fusion in CD13^KO^ cultures is not due to enhanced proliferation rates (**Fig. S1B)**. To circumvent the technical obstacle that BM-derived progenitor cells fuse poorly on the glass substrates required for high resolution optical immunohistochemical analysis, we adapted an alternate method of treating optical glass that enabled macrophage fusion (43). WT and CD13^KO^ thioglycollate-elicited peritoneal macrophages (TG-macs) seeded on plastic for Giemsa (**Fig 1B, E**) or treated optical glass for IF staining (**Fig 1C, F**) readily fused in the presence of MGC cytokines, IL-4+IL-13. This recapitulated our results in BM-derived progenitors as identified by brightfield and positive immunostaining for Mac-3, the mouse macrophage differentiation antigen, where the fusion index was significantly higher in the absence of CD13 (plastic; 2 fold and optical glass; 2.5 fold). Taken together, we conclude that CD13 acts as a physiologic regulator to control the rate of myeloid cell fusion.

**FIG 1.**
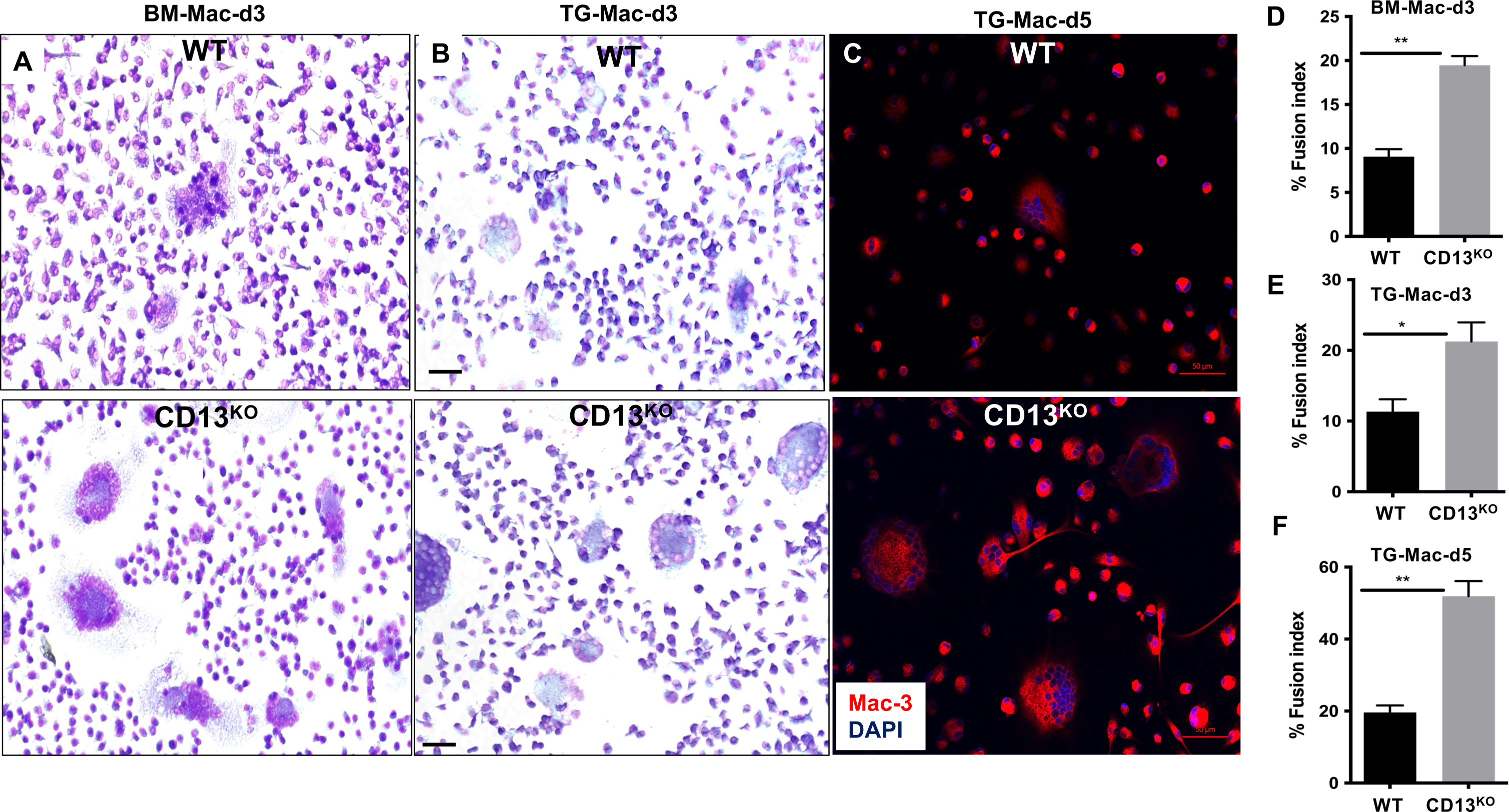
Lack of CD13 leads to exaggerated MGC formation in vitro. Giemsa staining of flow sorted, live (CD3/B220/Nk1.1)^-^ CD11b^lo/-^ CD115^hi^ Ly6G^+^ BM progenitor cells from WT and CD13^KO^ mice at d3 post MCSF+ IL-4+ IL-13 treatment. Cytoplasm; blue, Nuclei; pink/purple. **A)** BM-derived progenitor cells differentiated in presence of MCSF for d3 followed by IL-4+IL-13 for an additional 3d. **B)** Thioglycolate-elicited WT and CD13^KO^ peritoneal macrophages at d3 post IL-4+ IL-13 treatment on plastic. **C)** IF analysis of thioglycollate elicited WT and CD13^KO^ peritoneal macrophages at d5 post IL-4+ IL-13 treatment on treated optical glass surface with Mac-3^+^ (red) and DAPI; blue, nuclear stain. Magnification; 20x. **D-F**. % Fusion index of WT and CD13^KO^ BM derived MGC at d3, **D)** Thioglycollate-elicited MGC at d3 on plastic **E)** and Thioglycollate-elicited MGC at d5 on treated optical glass **F)** Cells with >3 nuclei indicate hypernucleated giant cells. Data represents +/-SD of 3 independent experiments. N=6/genotype, *;p<0.05, **;p<0.01. Scale bar; 50 μm.

### CD13 Promotes FBGC Formation and Fibrosis *in vivo*

Synthetic mesh implants are frequently used in hernia and other soft tissue repair and often induce macrophage fusion to form Foreign Body Giant Cells (FBGC) that are the histologic hallmark of the FBR. While their histological presence is characteristic of the FBR, it is unclear whether FBGCs directly contribute to the debilitating fibrosis of the FBR or are merely bystanders (8–11, 44), as abundant cytokines and chemokines, proteases, acids and free radicals are also present at the implant site (6, 45–47). To determine if CD13 contributes to FBGC formation *in vivo* and in turn, whether FBGCs themselves influence the FBR we used an *in vivo* mesh implant model where small sterile Polyethylene (PE) mesh implant was inserted subcutaneously into the back of 8-10 wk-old WT and CD13^KO^ mice of both genders. Implants were harvested at d1, 7 and 14 followed by evaluation of structural and inflammatory parameters in FFPE (Formalin-Fixed Paraffin Embedded) tissue implants. Immunohistochemical analysis indicated that the number of Mac-3^+^ FBGCs (activated macrophages with >3 nuclei indicated by white arrows) was significantly increased (>2-fold) in implants isolated from CD13^KO^ mice compared to WT counterparts at d14 (**Figs 2A-C, S2C)** supporting our *in vitro* MGC fusion data, with a concomitant increase the circulating levels of the cytokines IL-1β (1.8 fold) and TNF-α (2.2 fold) at d14 compared to d1 post implant (**Figs 2**D, E), indicating systemic inflammation. Of note, at d7 post-implant, neither the number of FBGCs nor levels of serum cytokines were significantly different between the genotypes. Furthermore, strong CD13 membrane expression was seen in the WT FBGCs (**Fig 2F,G**) and MGC (**Fig S2A-C**) and at the cell-cell contacts, consistent with a functional contribution of CD13 in FBGC and FBR.

**FIG 2.**
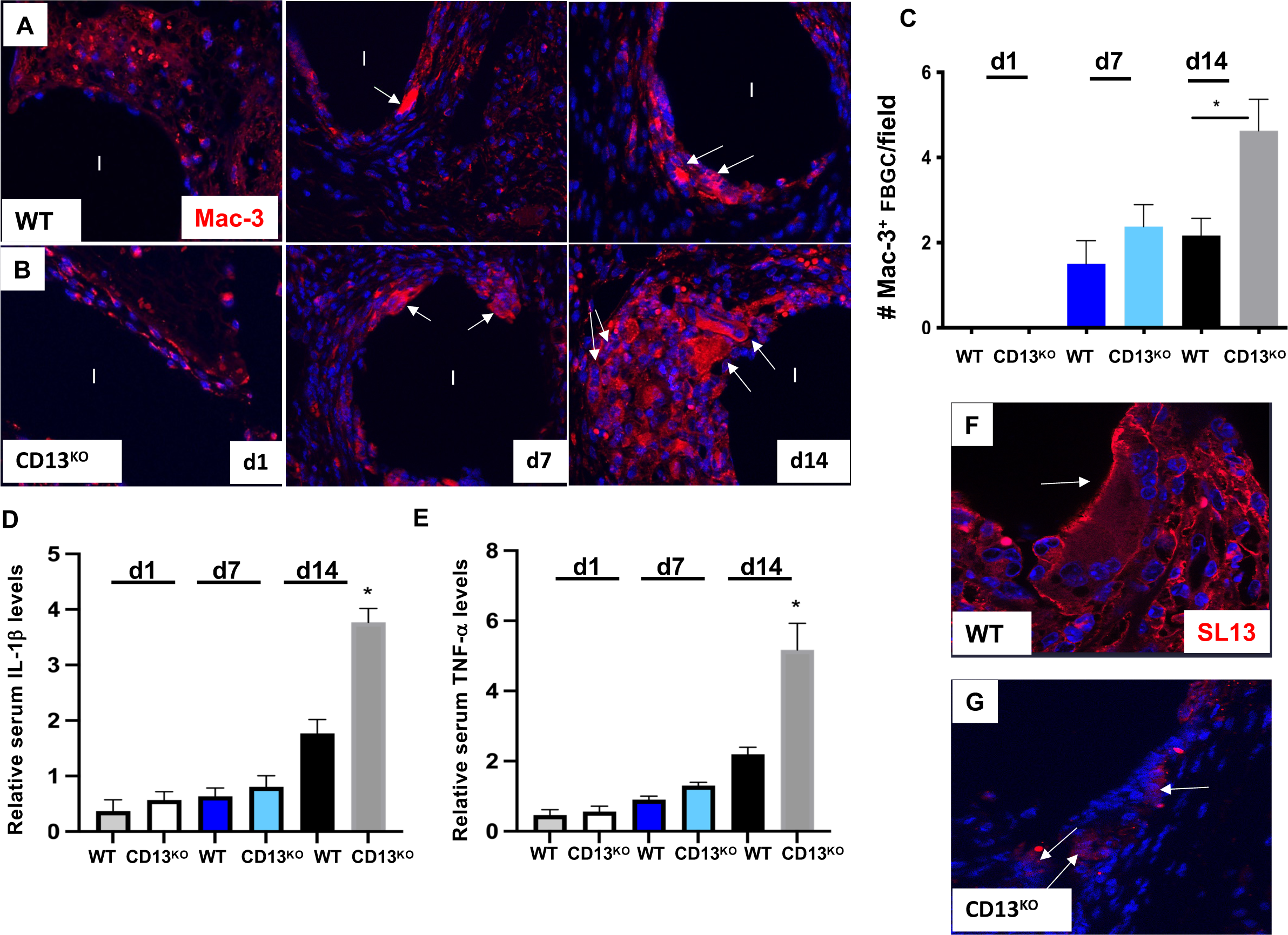
CD13^KO^ mice show amplified FBGC formation in an *in vivo* mesh implant model. **A-B.** Mac-3^+^ (red) Foreign body giant cells (FBGC with >3 nuclei) surrounding the mesh (indicated as I) in WT **A)** or CD13^KO^ **B)** mice at d1, 7 and 14 post-implant in a PE mesh implant model. **C)** Quantification of Mac-3^+^ FBGC per field over time indicate enhanced FBGCs in absence of CD13 at d14 post implant. **D-E)** Implants in CD13^KO^ mice induced elevated serum cytokine levels (IL-1β and TNF-α) compared to WT mice at d14 post implant. **F-G**. CD13 (SL13 mAb; red) is highly expressed in WT FBGC membrane **F)** but not in CD13^KO^ FBGC **G).** White arrows indicate FBGCs. Data represents +/- SD of 3 independent experiment. N=5/genotype, *;p<0.05. Scale bar; 50μm.

Collagen accumulation at the implant site is indicative of tissue fibrosis and advanced-stage FBR (47, 48). Image J analysis of Masson’s trichrome-stained implants showed a 1.89-fold increase in peri-implant collagen deposition in CD13^KO^ compared to WT at d14 post implantation (**Fig. 3A-C**), consistent with the notion that heightened numbers of FBGCs promote the evolution to fibrosis. At the earlier 7d timepoint, fibrosis was mild and not significantly different between genotypes. In agreement with enhanced fibrosis, IF analysis of TGFβ and quantification by Image J analysis in implants showed that the intensity of pro-fibrotic TGFβ expression in the FBGCs themselves was 1.84-fold higher at CD13^KO^ implant sites compared to WT counterpart (**Fig. 3D, E)** although importantly, the number of F480^+^ macrophages remained equivalent between genotypes (**Fig. 3F, G)**. We conclude that the CD13-dependent increase in the number of mesh-induced, peri-implant FBGCs exacerbates fibrosis and implicates a direct impact of FBGC numbers on FBR progression.

**FIG 3.**
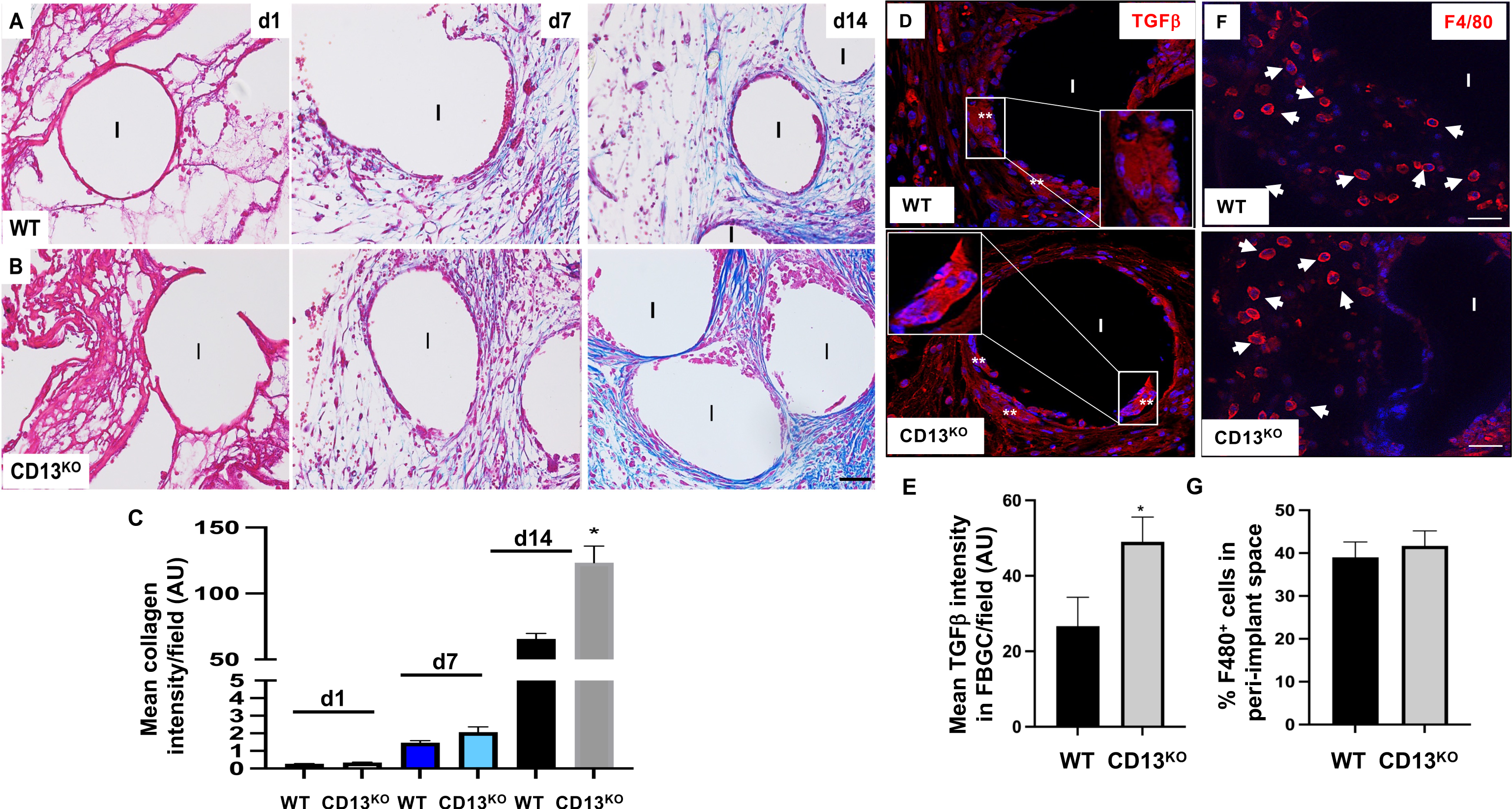
FBGC-induced fibrosis is accelerated in CD13^KO^ an *in vivo* mesh implant model. Masson’s trichrome staining indicates enhanced peri-implant (implant; I) collagen deposition (blue, fibrosis) in CD13^KO^ mice **B**) compared to WT **A**) at d14 but not at d1 or d7 post implant. Quantification of the trichrome-stained collagen content **C**) per field over time by Fiji analysis. **D,E)** Increased TGFβ expression in CD13^KO^ FBGCs (double white asterisks) at the site of the implant compared to WT as indicated by the intensity of TGFβ in FBGCs/field analyzed by Fiji software. **F,G)**. IF analysis of implants indicated equivalent numbers of F480^+^ macrophages (red, indicated by white arrows) in the peri-implant region in both genotypes. Magnification; 20x. Scale bar; 50μm. *; p<0.05. Data represents +/- SD of 3 independent experiment. N=3/genotype.

### CD13 controls MGC fusion by regulating actin protrusion and endocytic vesicle formation

In addition to the fusogenic proteins, cytoskeletal organization and rearrangements are also crucial for MGC formation. Studies have shown that actin filaments are involved in the formation of fusion pores, which allow the exchange of cytoplasmic contents, necessary for the cells to merge and form a single cell (22, 49–53). In addition, the processes of receptor endocytosis/recycling and the formation of membrane protrusions, are key mechanisms controlling cell-cell fusion (3, 19, 43, 49, 50, 54) as demonstrated by the abundant actin^+^ cell extensions formed in MGCs in response to IL-4-induction (4, 55) and abrogation of macrophage fusion by actin depolymerization (56, 57). Previously, we and others have shown that CD13’s intracellular domain localizes and tethers signaling molecules to cytoskeletal proteins at the membrane during receptor recycling, suggesting that CD13 may directly impact cytoskeletal structures such as actin protrusions (33, 35, 38, 58). Giemsa staining of BM progenitors at the early stages of MGC fusion induction showed that CD13^KO^ macrophages produced significantly more (2.8 fold; **Fig. 4A-C)** and longer (1.7 fold; **Fig. S3A**) actin protrusions compared to WT cells as early as d2 post-induction, linking CD13 expression to the formation of actin-based protrusions as we had previously demonstrated in endothelial cells (59). Interestingly, mononuclear cells appear to attach to MGCs along the length of the protrusions more frequently in CD13^KO^ as compared to WT cells (**Fig. S3B**) indicated by Phalloidin staining, which may provide clues as to why fusion accelerates under conditions supporting actin protrusions. To further establish that these CD13-dependent actin protrusions are responsible for the hyperfusion phenotype, we inhibited actin polymerization with Cytochalasin D at d1 post-cytokine stimulation and saw significantly reduced MGC formation indicated by ∼4 fold reduction in the fusion index in CD13^KO^ myeloid progenitors compared to vehicle, linking actin dynamics to the hyperfusion phenotype (**Fig. S3C**).

**FIG 4.**
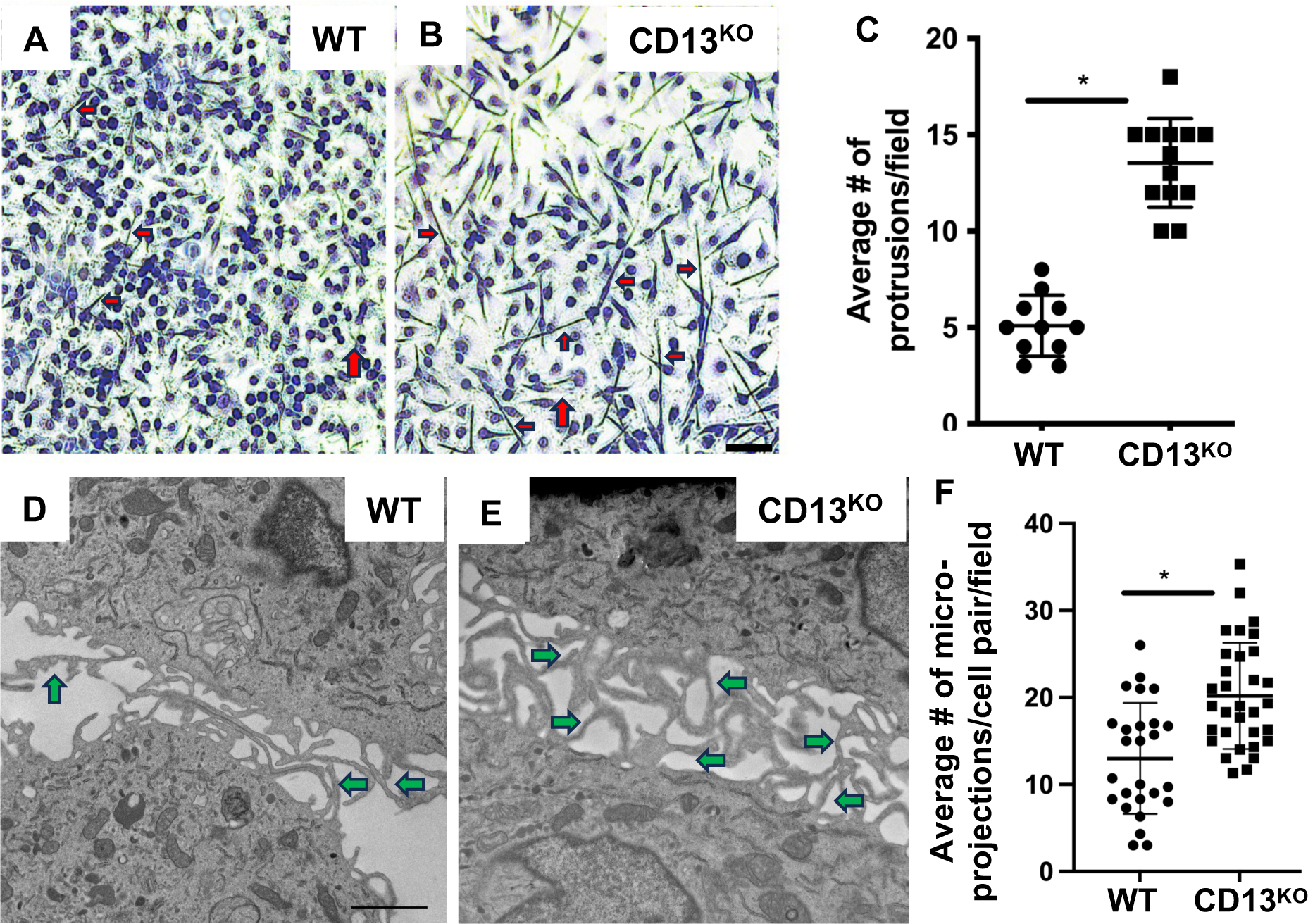
Actin cytoskeletal changes at the pre-fusion stage drives MGC formation in the absence of CD13. **A-B)** Brightfield images of Giemsa stained WT and CD13^KO^ flow-sorted BM macrophages grown on plastic indicate an increased number of actin protrusions at d2 post MCSF+ IL-4 + IL-13 treatment in CD13^KO^ **B**) compared to WT **A**) cells. Red arrows indicate protrusions. **C**) Average number of protrusions is significantly higher in absence of CD13 at d2 post MCSF+ IL-4 + IL-13 treatment. **D-F**. Transmission Electron Microscopy of WT **D**) and CD13^KO^ **E**) flow sorted BM macrophages at d2 post MCSF+IL-4+IL-13 treatment. Green arrows indicate microprojections between two WT **D**) and CD13^KO^ **E**) neighboring cells . **F**) Quantification of microprojections/cell pair ∼1μm apart. Data represents average of 2 isolates and ∼26-33 fields/isolate evaluated by three individuals in a blinded manner. Data represents +/- SD of 2 independent experiments. N=3/genotype. *;p<0.05. Scale bar; A-B; 50μm, D-E; 5μm.

To explore the cell cytoskeletal changes required for proper cell-cell fusion at the organelle-scale, we performed high resolution Transmission EM (TEM) to determine potential differences in topography, morphology and arrangement of cellular protrusions (3, 43). TEM has been previously used to study osteoclast, but not MGC fusion (refs). Since our standard growth and fusion conditions in 96-well dishes were challenging for TEM processing, we developed a method to grow MGCs on Aclar plastic (5mm^2^, suitable for a 96-well dishes) and found that macrophages readily fused under these conditions, enabling EM analysis of sites of cell-cell contact for individual cell protrusions (60–62). Ultrastructural examination of fusing WT or CD13^KO^ myeloid cells at d2 post fusion revealed microprojection-like structures that join two neighboring cells, not visible under brightfield microscope (≥1μm apart, indicated by green arrows, **Fig 4F, G**). The total number of micro-projections represents both fused and non-fused microprojections between two neighboring cells. A total of 26 WT images and 33 CD13KO images were counted and each point represents average count measured by three separate individuals in a blinded manner. Analysis of EM images indicated a significant increase in micro-projections between fusing CD13^KO^ cells compared to the WT counterparts (**Fig. 4H**), supporting our notion that CD13 impacts cytoskeletal dynamics and protrusion formation during fusion.

### CD13 mediates fusogen expression by post-transcriptional mechanisms

The “master fusogen” DCSTAMP and the tetraspanins CD9 and CD81 are common regulators of cell-cell fusion whose expression is transiently induced upon stimulation of MGC fusion, but must be subsequently reduced to allow fusion to proceed (3, 16, 19, 21, 23, 63–65). Importantly, the sustained expression of fusogens has been shown to accelerate fusion both *in vivo* and *in vitro* (28, 39, 66). Indeed, prolonged CD81 expression has been shown to lead to abnormal fusion and MGC formation (67). To monitor fusogen trafficking, we tracked the loss and surface re-expression (recycling) of fusogens in WT and CD13^KO^ macrophages under fusogenic conditions by surface biotinylation and capture ELISA as described previously (38). Briefly, at d5 post-fusion induction, proteins on the surface of TG-elicited peritoneal macrophages from WT and CD13^KO^ mice were surface biotinylated and allowed to internalize at 37° for 30 min, followed by chase in complete medium over time to allow labeled proteins to recycle to the surface. At each time point of cell harvest, surface biotin was removed by MesNa treatment to eliminate the recycled fusogens, allowing us to measure biotinylated internalized fusogen levels. Next, cell lysates were incubated in wells coated with anti-CD9 or anti-DCSTAMP Ab and captured biotinylated fusogen was measured by secondary mAb detection. The relative proportion of recycled CD9 and DCSTAMP at the surface was determined by the relative amount of internalized fusogen in cells during chase. Fusogen recycling was calculated by assuming that the sum of labeled internalized + recycled proteins = 100% and that the degree of recycling of the fusogen to the surface at each time point during chase = 100% minus the % of the labeled fusogen remaining inside the cell at each time point. In agreement with the sustained fusogen surface expression in CD13^KO^ osteoclasts (39), fusogen surface expression over time was significantly enhanced on CD13^KO^ cells (1.5-1.8 fold), suggesting that CD13 dictates the levels of these key fusion regulatory molecules (**Fig. 5A,B**). Furthermore, the transcript levels of these fusion regulators were equivalent in WT and CD13^KO^ fusion cultures (**Fig. S4**), indicating that CD13 regulated fusogens by post-transcriptional mechanisms. To further verify this concept, we blocked *de novo* protein synthesis in WT and CD13^KO^ TG-macs with cycloheximide, followed by treatment with MG-132 to inhibit both the proteasomal and lysosomal degradation pathways (68). Quantification of surface expression by flow cytometry indicated that blocking protein synthesis for 6h reduced the abundance of CD9 and DCSTAMP proteins on the surface to a greater extent in WT compared to CD13^KO^,suggesting that lack of CD13 stabilizes surface expression of DCSTAMP and CD9 by controlling protein turnover via proteasomal and lysosomal degradation pathways (**Fig. 5C-D, S5**).

**FIG 5.**
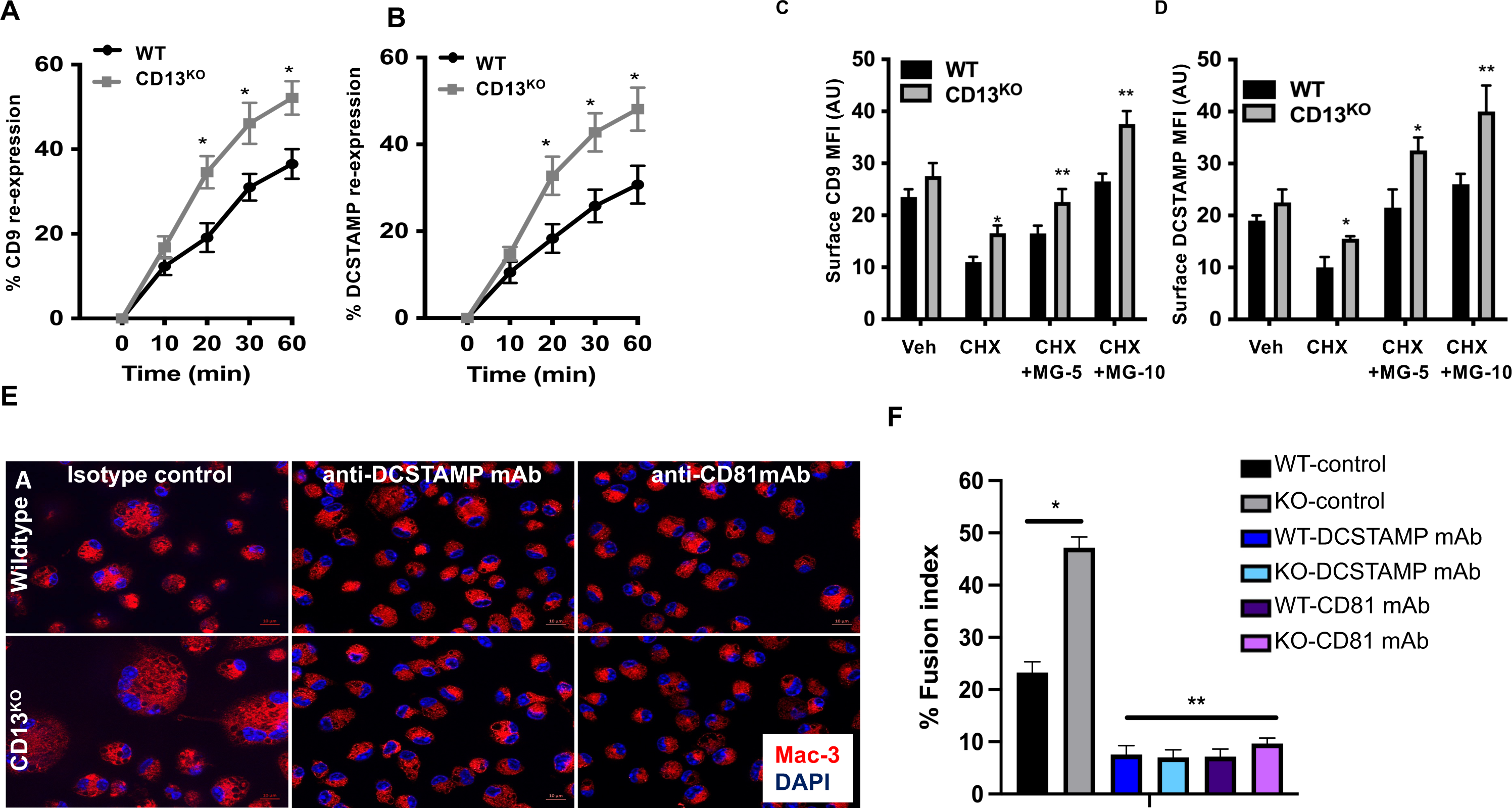
CD13 mediates protein stability via a ubiquitination pathway. **A-B)** Surface expression of biotinylated CD9 **A**) and DCSTAMP **B**) measured by capture ELISA with anti-CD9 and anti-DCSTAMP Abs indicate % surface fusogen expression over time is higher in CD13^KO^ compared to WT TG-peritoneal macrophages at d3 post treatment with MCSF+ IL-4 + IL-13. **C-D)** Flow-sorted BM-myeloid progenitors under fusogenic conditions were treated with cycloheximide, (CHX; protein synthesis inhibitor) (100μg/ml) +/- 5 & 10μM MG-132, (degradation inhibitor) for 8h. Flow cytometric analysis of surface mean fluorescence intensity indicated a rapid decay of CD9 and DCSTAMP in WT compared to CD13^KO^ in presence of CHX vs. CHX+ MG132. **E-F**) Functionally blocking DCSTAMP and CD81 fusogens rescues abnormal fusion phenotype in CD13^KO^ indicated by % fusion index. TG-WT and CD13^KO^ peritoneal macrophages treated with blocking DCSTAMP and CD81 mAbs (20μg/ml) in presence of IL-4 + IL-13 at d1 were allowed to fuse for d7 followed by immunohistochemical analysis **E**) and measurement of fusion index **F**). IF analysis of Mac-3^+^ cells (red) showed a significant reduction in MGC formation in CD13^KO^ similar to the level of the WT cells by d7 post treatment. Data represents +/- SD of 3 independent experiments. N=3/genotype. **;p<0.01, *;p<0.05. Scale bar; 10μm.

To establish a direct link between CD13-dependent fusogen expression and fusion, we treated WT and CD13^KO^ peritoneal macrophages with DCSTAMP and CD81 blocking mAbs at 20μg/ml daily for 7d post IL-4+IL-13 addition (67, 69, 70). IF analysis of Mac-3^+^ MGCs revealed that in comparison to isotype control, blocking the fusion-promoting proteins clearly reversed the hyperfusion phenotype in CD13^KO^ cells to a level similar to WT at d7 post cytokine stimulation (**Fig. 5E-F**). Together, these results confirm that manipulation of fusogens by either fusogen blocking mAbs or lack of CD13 correlates with the hyperfusion phenotype.

### CD13 regulates ubiquitination of fusogens

While little is known about the post-translational fate and regulation of the fusogens in giant cell formation, it is thought that ubiquitination and degradation of fusion proteins is critical to ensure proper cell-cell fusion into multinucleated cells (71, 72). This process involves the tagging of fusion proteins with ubiquitin, a small protein that marks proteins for degradation by the proteasome (73). Various studies have shown that ubiquitination is a multi-step, tightly controlled enzymatic process and when dysregulated, leads to abnormal cell function including fusion and multinucleated cell formation (74–77). Therefore, to determine if CD13 participates in the ubiquitination pathway, WT and CD13^KO^ peritoneal macrophages treated with different doses of the degradation inhibitor MG-132 for 3h were lysed and incubated on a PROTAC assay plate (LifeSensors) for 2h to capture polyubiquitinated proteins. Bound, polyubiquitinated DCSTAMP or CD9 protein was detected by ELISA which were significantly enhanced (2-fold) in WT cells, resulting in decay of the fusogens compared to CD13^KO^ cells (**Fig. 6A, B**), indicating that CD13 controls protein levels via a ubiquitination pathway.

**FIG 6.**
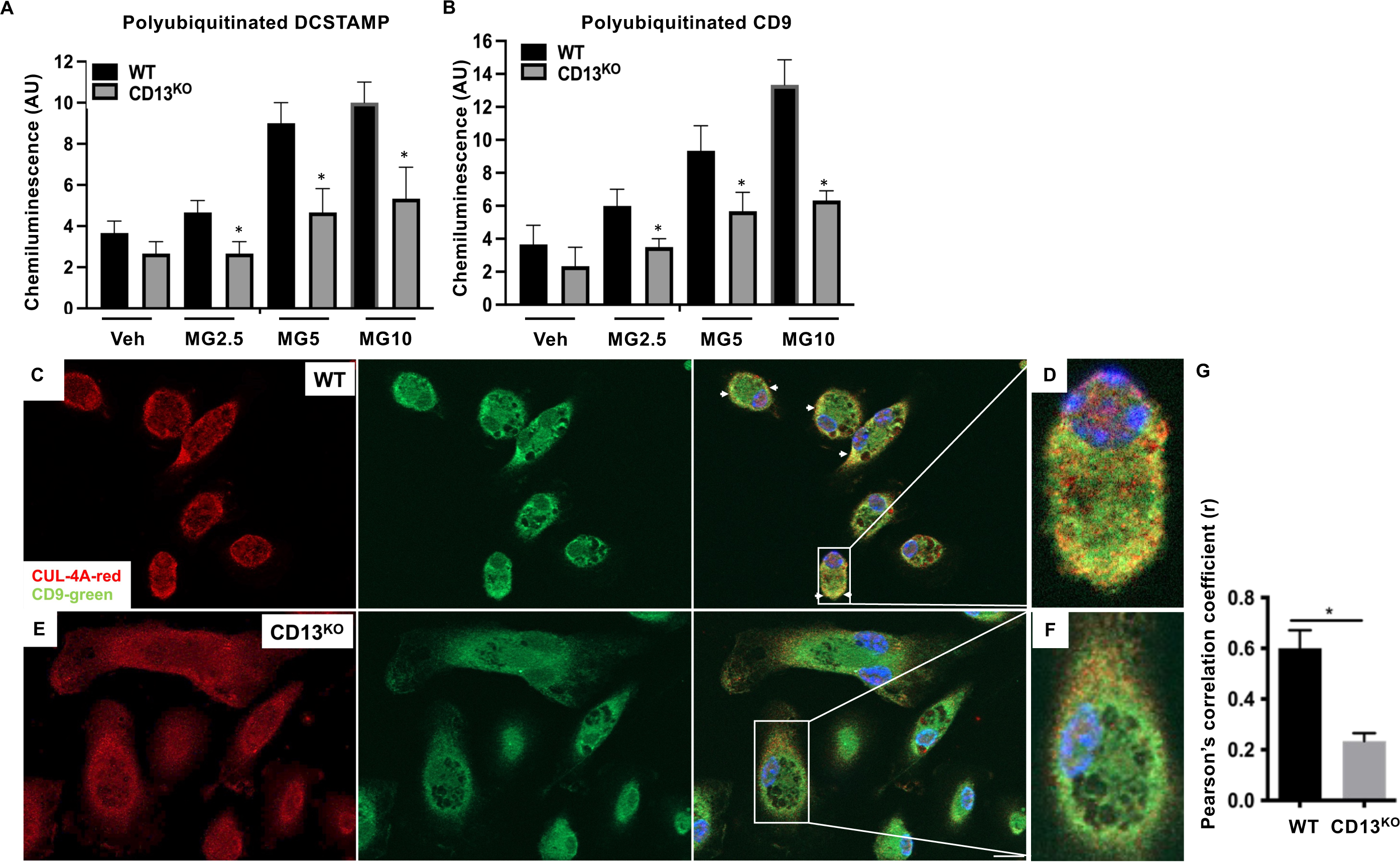
Loss of CD13 results in reduced ubiquitination of fusogen and mislocalization of Cullin-4A and CD9. **A-B.** WT and CD13^KO^ TG macrophages were treated with increased doses of proteasomal degradation inhibitor MG-132 for 0-3h. Cells were lysed & subjected to ELISA-based protein ubiquitination assay using a PROTAC assay plate containing polyUb capture reagent. Bound, polyubiquitinated target proteins were detected with anti-DCSTAMP **A**) or CD9 **B**) mAb. Chemiluminescence analysis indicated reduced ubiquitination of fusogens in CD13^KO^ compared to WT cells. **C-E**. Cullin-4A and CD9 are mislocalized in CD13^KO^ compared to WT TG-macrophages at d3 under fusogenic conditions. Cytoplasmic expression of CUL-4A (red) in WT **C**) is markedly altered in CD13^KO^ **E**) myeloid cells undergoing fusion. **D-F**) Zoomed image of colocalization of CUL-4A (red) and fusogen CD9 (green) in the WT (arrows) but not in the CD13^KO^ cells, further confirmed by Pearson’s correlation coefficient (r) **G**). Magnification; 63x oil. DAPI; blue. Data represents +/- SD of 3 independent experiments. N=3/genotype. *; p<0.05. Scale bar; 10µM.

### CD13 is required for the association and localization of Cullin-4A and fusogens

In unpublished mass spectrometry studies, we had previously determined that the scaffold protein Cullin 4A (CUL-4A) co-immunoprecipitated in a complex with CD13 (not shown). CUL4 is a core component of the CRL4 E3 (Cullin Ring Ubiquitin E3) ligase complex which is responsible for the recruitment, ubiquitination and degradation of numerous proteins (78). Defects in the expression and/or localization of CUL4 protein have been shown to compromise ubiquitination and subsequent target protein turnover (79–81). As a scaffold, CD13 organizes molecular complexes by bringing key proteins into close proximity to promote their interactions and situating these complexes at the proper location for optimal function. Therefore, lack of CD13 could disrupt the interaction and localization and of key complexes (82). IF analysis to examine potential effects of CD13 on CUL-4A, CD9 and DCSTAMP localization during fusion of WT and CD13^KO^ TG-elicited peritoneal macrophages indicated that in contrast to the primarily juxtamembrane expression in WT cells, the expression pattern of CUL-4A is more diffuse throughout the cytoplasm in the absence of CD13 (**Fig. 6C, E**). Importantly, colocalization of CUL-4A and CD9 (Pearson’s correlation coefficient, WT vs. CD13^KO^; r=0.7 vs. 0.3) and CUL-4A and DCSTAMP (Pearson’s correlation coefficient, WT vs. CD13^KO^; r=0.52 vs. 0.2) was significantly reduced in CD13^KO^ cells as compared to WT (**Fig. 6D, F, S6**), in agreement with CD13 acting as a scaffold to promote the association of the fusogenic proteins and CUL-4A for proper ubiquitination and degradation.

### CD13, ubiquitination and actin binding protein transcripts are enriched in FBGCs

In a recent study, sc-RNAseq analysis of cells at sites of mesh-induced FBR showed that Actn1 [α-actinin, CD13-interacting cytoskeletal protein, (35)], Dcstamp (fusogen) and Uchl1 (ubiquitin C-terminal hydrolase) are highly expressed in FBGCs compared to M2 macrophages or precursors (51). Importantly, further mining of the RNAseq data revealed that Anpep (CD13) is highly expressed in giant cells compared to the precursor populations (**Fig. S7),** together supporting our results that mechanisms regulating ubiquitination and cytoskeletal rearrangement pathways are important in FBGC formation. Taken together, we conclude that CD13 is a critical regulator of myeloid cell fusion, both in the formation of MGCs *in vitro* and FBGC formation *in vivo* (this study) as well as in osteoclast fusion *in vitro* and *in vivo* (39). Mechanistically, CD13 limits formation of microprojections and actin-based protrusions at the pre-fusion stage, while facilitating the association of fusogens with the CRL4 complex during fusion. Therefore, loss of CD13 leads to hyperfusion, exaggerated microprojections between neighboring cells, compromised ubiquitination and subsequent degradation of fusogens by the CRL4 complex, contributing to accelerated fusion (**Fig. 7)**. Together, our results suggest that CD13 may be a therapeutic target to control the damaging inflammation observed at sites of implantation and prevent implant failure.

**FIG 7.**
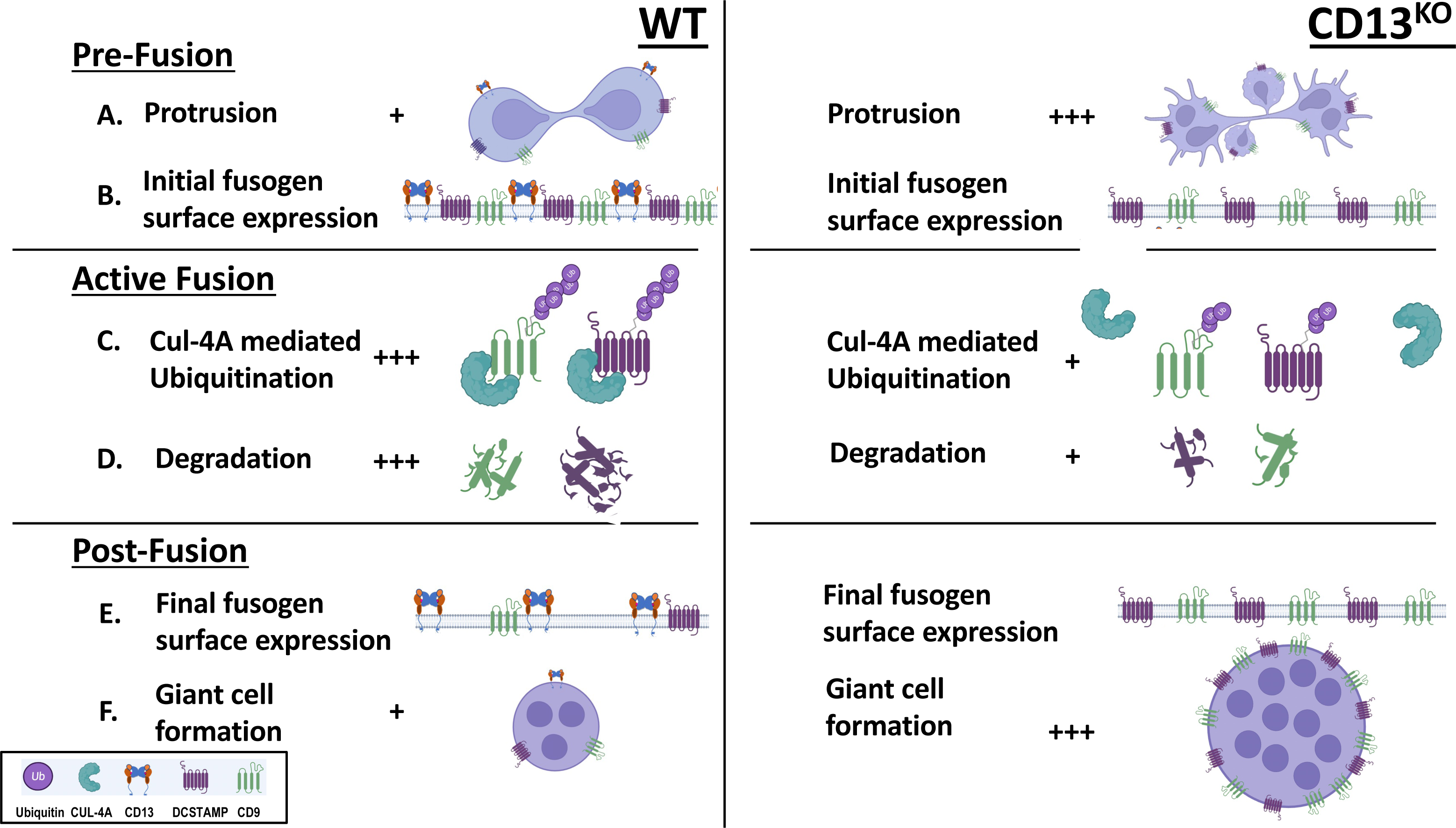
Schematic of CD13-mediated giant cell fusion by a post transcriptional regulation of fusogens. Loss of CD13 results in increased protrusion and microprojection formation between neighboring cells but comparable fusogen expression among genotypes at the pre-fusion stage **A,B)** with reduced fusogen ubiquitination/degradation during fusion, augmenting fusogen surface expression **E)** and ultimately, multinucleated giant cell formation **F).**

## Methods

Global young (8–10 weeks) wildtype and CD13^KO^ (C57BL/6J) male and female mice were generated and housed at the Gene Targeting and Transgenic Facility at University of Connecticut School of Medicine. All procedures were performed in accordance with the guidelines and regulations approved by the University of Connecticut School of Medicine Institutional Animal Care and Use Committee. University of Connecticut School of Medicine is fully accredited by Association for Assessment and Accreditation of Laboratory Animal Care (AAALAC) International and Public Health Service (PHS) assurance number is A3471-01 (D16-00295) and USDA Registration Number is 16-R-0025.

Euthanasia by CO2 followed by cervical dislocation was performed and is an accepted method consistent with AVMA Guidelines for the Euthanasia of Animals to minimize the pain or discomfort in animals.

### Flow Cytometry

BM progenitors were stained with the following antibodies for phenotypic analysis and sorting: Anti-CD3 APC (Biolegend 100235), anti-B220 APC (Biolegend 103212), anti-NK1.1 APC (Biolegend 108710), anti-CD11b e450 (Invitrogen 48-0112-82), anti-CD115 PE/Cy7 (Biolegend135524), anti-Ly6C AF700 (Biolegend 128024), and UV Live dead dye (Invitrogen L23105). Flow cytometry for live cells derived from WT and CD13^KO^ bone marrow were performed with BD FACS aria to obtain (CD3/B220/Nk1.1)^-^ CD11b^lo/-^ CD115^hi^ Ly6G^+^ progenitor cells. Data was analyzed using BD FACS DIVA version 9.0 and FlowJo 9.9 version software. For measurement of surface fusogens, anti CD9 antibody-FITC (Biolegend; clone MZ3) and anti-DCSTAMP antibody (Creative Diagnostics; Clone 2B3) were used.

### Isolation and culture of macrophage progenitors from bone marrow

BM cells were obtained by isolating the femur and tibia from WT and CD13^KO^ mice of both genders. The ends of the femur and tibia are removed to expose the bone marrow (BM). BM cells were harvested by flushing through the BM with 10 ml 1x PBS supplemented with 2% fetal bovine serum. Cells were treated with RBC lysis buffer and filtered through a 40 μm cell strainer (Corning 431750). Flow cytometry was used to isolate (CD3/B220/Nk1.1)^-^ CD11b^lo/-^ CD115^hi^ Ly6G^+^ progenitor cells. Progenitor macrophages were seeded at density 20,000-25,000 cells per well on 96 well dish on d0 in DMEM (Gibco 11965092) with 10% FBS and 1% penicillin-Streptomycin. Cells were treated with 20 ng/mL M-CSF for 2d for macrophage differentiation and proliferation, followed by treatment with 30 ng/mL each Il-4 (R&D Systems; 404ML010) and IL-13 (R&D Systems; ML010) cytokines in fresh complete media and incubated at 37 degree C with 5% CO_2_ for an additional 3-5d to allow fusion to proceed.

### Generation and culture of Thioglycollate-elicited macrophages from peritoneum

Both WT and CD13^KO^ mice of both genders were intraperitoneally injected with 1 mL 4% Brewers Thioglycollate (TG) solution (BD; 211716). After 48 hours mice were then euthanized via CO2 and cervical dislocation and injected with 10 mL cold PBS into the peritoneal cavity to collect peritoneal macrophages. Peritoneal lavage cells suspended in PBS were extracted and treated with RBC lysis buffer for 5 min. Peritoneal Lavage cells were centrifuged at 1200 rpm for 5 min and resuspended in alpha MEM (Gibco; 12571063) media supplemented with 10% fetal bovine serum (Gibco; 16250078) and 1% penicillin-streptomycin (Gibco; 15140122). Total live cells counted manually with Bright Line Hemacytometer with Trypan Blue dye. 50,000-75,000 cells per well were seeded on a 24 well dish on d0. Fusion inducing cytokines IL-4 (R&D Systems; 404ML010) and IL-13 (R&D Systems; ML010) were added in 60 ng/mL each in complete media on d3 and incubated at 37 degree C with 5% CO_2_ for an additional 3-5d to allow fusion to proceed. **In Vivo Implant Analysis**

WT and CD13^KO^ mice of both genders were anesthetized with isoflurane. Prior to implantation, mice were injected subcutaneously with 8mg/kg Bupivacaine (Hospira NDC; 0409-1163-18) at the site of implantation to numb the area. A 0.5 inch × 0.5 inch sterile polypropylene mesh implant (Ethicon; PMXS) was inserted subcutaneously in the flank region of the mice on d0 followed by peritoneal injection with 30 µL Meloxicam (Covetrus NDC; 11695-6936-1) to reduce pain. Implants were harvested on d1, d7, and d14 following euthanasia with CO_2_ and cervical dislocation. Harvested implants were fixed in 4% formalin overnight and embedded in paraffin. FFPE (Formalin-Fixed Paraffin Embedded) implant sections were cut via microtomy into 5 µm sections and loaded onto microscope slides. To process for imaging, slides were deparaffinized via submersion twice in xylene for 10 min each, submerged in 100% and 70% ethanol for 5 min each, and washed with water for 5 min at room temp. Slides were then subject to immunofluorescence or light microscopy as indicated.

### Preparation of optical glass for TG-elicited macrophage fusion

Optical cover glass was acid cleaned with 12N HCl, washed with deionized water, rinsed with ethanol and allowed to dry for overnight. Acid-cleaned cover glass was treated with 1mg/ml Paraffin wax (SIGMA) solubilized in toluene for few mins, excess solution was removed, washed with sterile water, allowed to dry and sterilized under UV light.

### Immunofluorescence assay

Thioglycollate-elicited peritoneal macrophages (TG-Macs) grown on coverslips were washed twice with 1x PBS and fixed with 4% Paraformaldehyde for 20 min. Cells were then permeabilized for 5 min with 0.01% Triton-X and blocked for an hour with 5% BSA. Primary antibodies (see list above) were added in 1% serum in 5% BSA buffer and incubated at 4°C overnight. Coverslips were washed twice with 1x PBS and Alexa Fluor secondary antibodies were added in addition to DAPI (nuclear stain) in 1% BSA and incubated at 4°C overnight. Coverslips were then washed with PBS and mounted onto microscope slides with Prolong Gold Antifade mounting media (1 drop, Molecular Probes; P10144).

For tissue sections, post-deparaffinization, implant sections were subjected to antigen retrieval with 10 mM sodium citrate and microwaved for 1 min and 25 sec. Slides were blocked with 5% serum in 5% BSA buffer for 1 hr at room temperature. Slides were washed twice with 1x PBS and primary antibodies were added with 1% Serum in 5% BSA buffer and incubated at 4°C overnight. Alexa Fluor secondary antibodies and DAPI were added in 1% BSA and incubated away from light for 1 hr at RT. Coverslips were added to slides with the addition of Prolong Gold Antifade mounting media (1 drop, Molecular Probes; P10144). All samples were imaged at excitation wavelength of 488 nm (Alexa 488), 543 nm (Alexa 594) and 405 nm (DAPI). Samples were imaged by Zeiss LSM 880 confocal fluorescence microscope and analyzed by using Zeiss Zen 2.0 Pro blue edition software

### Giemsa Staining

BM progenitors or TG-macs grown on plastic were fixed with 100% methanol for 10 min. After fixation, methanol was aspirated, and cells were left to air dry for at least one hour. Cells were then stained with Giemsa (Harleco; R03055) diluted 1:20 in dH_2_O for 25 min. Cells were then washed twice with dH_2_O and stored at room temp away from light. Cells were imaged via light microscopy using Zeiss fluorescence inverted microscope and analyzed by using Zeiss Zen 2.0 Pro blue edition software. Average protrusion length of fusing cells was measured with ImageJ software (https://imagej.nih.gov/ij/).

### Measurement of Fusion index

Random fields with similar cell densities of Giemsa stained or immunofluorescence images were considered for evaluation of fusion index and quantified in a blinded manner. % Fusion index= [Total number of nuclei in fused cells (>3) / Total number of nuclei in fused and non-fused cells] × 100.

### Masson Trichrome Staining

Post deparaffinization, in vivo implant tissue was stained with Masson trichrome. Masson Trichrome staining was performed at the Research Histology Core Labs at UConn Health. Intensity of fibrosis by Masson Trichrome was calculated using ImageJ software (https://imagej.nih.gov/ij/).

### Electron Microscopy

Flow sorted mouse bone marrow cells were cultured in 96-well dishes as described above. To make cells amenable to processing for electron microscopy, pieces of Aclar plastic (Electron microscopy sciences; 50425-10) were placed in the bottom of the 96 well dishes, allowing cells to grow and fuse on the removable substrate.

Post fusion, the cells grown on Aclar were fixed in 2.5% glutaraldehyde (EMS) in 0.1M sodium cacodylate buffer (EMS) on ice for 15-30 min. Cells were rinsed in 0.1M cacodylate buffer five times, then post-fixed in 1% Osmium tetroxide (EMS) and 0.8% potassium ferricyanide in 0.1M cacodylate buffer for 15-30 min. Cells were then rinsed with Milli-Q filtered water and stained with 1% aqueous uranyl acetate (EMS) for 15-30 min. Samples were rinsed five times with water, then dehydrated in a graded series of ethanol (50%, 75%, 95%, 100%,) for 5-10 min each. Using an aluminum dish, the Aclar pieces with cells were further dehydrated in two changes of propylene oxide (EMS) for 2-3 min each. Next, the cells were infiltrated in Polybed 812 epoxy resin with BDMA in graded steps with propylene oxide. The cells on Aclar were flat-embedded between larger pieces of Aclar plastic with spacers made by layers of parafilm.

### TEM Image Acquisition/Analysis

Seventy nanometer sections were collected on copper mesh EM grids (EMS). Fusing WT and CD13^KO^ monocytes/macrophages that were within two micrometers from each other were imaged at 4000x on a Hitachi H-7650 at 80 kV. Cell protrusions (700x magnification) and microprojections (4000x magnification) between two neighboring monocytes at d2 post fusion that were at least 1mm apart were counted in a blinded manner.

### Quantitative RT-PCR

Total RNA was extracted using TRIZOL reagent (Invitrogen) according to manufacturer’s instruction and as described before{Ghosh, 2021}. Relative transcript level was normalized to GAPDH level. GenBank primer sequences (http://pga.mgh.harvard.edu/primerbank/) was used to determine fusogen primer sequences. PCR primer sequences used are as follows: *DC-STAMP*, 5′-GGGGACTTATGTGTTTCCACG-3′ (forward) and 5′-ACAAAGCAACAGACTCCCAAAT-3′ (reverse); *CD9*, 5′-ATGCCGGTCAAAGGAGGTAG-3′ (forward) and 5′-GCCATAGTCCAATAGCAAGCA-3′ (reverse); *CD81*, 5′-CAGATCGCCAAGGATGTGAAG-3′ (forward) and 5′-GCCACAACAGTTGAGCGTCT-3′ (reverse); *GAPDH*, 5′-GGATTTGGTCGTATTGGG-3′ (forward), 5′-GGAAGATGGTGATGGGATT-3′ (reverse) Transcript data was analyzed using CFX Manager version 3.1 (https://www.bio-rad.com/en-us/sku/1845000-cfx-manager-software?ID=1845000) (Biorad).

### Capture ELISA based Pulse-Chase Experiment

To assess re-expression of fusogens in cells undergoing fusion, WT or CD13^KO^ TG-macrophages at d5 were surface-labeled with 0.2mg/ml NHS-S-Biotin (Pierce) at 4°C for 30 min. Labeled cells were washed in cold PBS and internalization was allowed to proceed at 37°C for 30 min, at which point the remaining surface biotin was removed by treatment with 20mM MesNa in 50mM Tris-HCl (pH8.6) and 100mM NaCl for 15 min at 4°C. MesNa was quenched by lodoacetamide (20mM) treatment. Cells were then chased in label free complete medium for 30 min at 37°C. A second round of MesNa was administered to remove surface biotin from recycled fusogens to measure the relative amount of internalized fusogen in the cell lysate by capture ELISA. Cells were lysed with a protease inhibitor cocktail in RIPA lysis buffer. To quantify the amount of remaining biotinylated fusogens present in the cell lysate, 96-well microtiter plates were coated with 5μg/ml anti-CD9 (MyBiosource; MBS7113510) or anti-DCSTAMP (Creative Diagnostics; Clone 2B3) in 0.05M sodium carbonate (pH9.6) at 4°C. Plates were blocked in PBS containing 5% BSA at room temperature for 1 hour. Captured fusogens were measured by overnight incubation of cell lysate at 4°C. Plates were washed in PBS/Tween-20 and incubated with streptavidin-HRP for 1 hour at 4°C and biotinylated CD9 or DCSTAMP was measured by addition of a chromogenic agent ortho-phenylenediamine. Fusogen surface re-expression was calculated based on the assumption that % internalization + % surface re-expression = 100%, therefore % fusogen re-expression = 100% -% fusogens internalized present post chase. To assess fusogen re-expression without protein synthesis, the cells were treated with 100 µg/mL cycloheximide (CHX) for 6 hours prior to internalization. To further assess the role of fusogen recycling, the cells were treated with 100 µg/mL CHX and 5-10 µM MG132 for 6 hours prior to internalization.

### Chemiluminescence-based assay of polyubiquitination of target fusogens

WT and CD13^KO^ peritoneal macrophages under fusogenic condition were treated with MG132 at 2.5 µg, 5.0 µg and 10µg for 3h, followed by lysis with RIPA lysis buffer. 20 µg of cell lysate were added to PROTAC assay plate (LifeSensors; PA950) and incubated for at RT for 2h. Wells were washed and anti-DCSTAMP (Creative Diagnostics; Clone 2B3) or CD9 (MyBiosource; MBS7113510) Ab were added in 1x blocking agent and incubated at RT for 1h followed by addition of HRP-conjugated secondary Ab and incubated at RT for 1h. Detection reagents (supplied in the kit) were added and the plates were subsequently read by a microplate reader according to manufacturer’s instruction.

### Statistical analysis

Statistical analysis was performed using unpaired, two-tailed Student’s t test using GraphPad Prism software and results are representative of mean ± SD or +/- SEM as indicated. Differences at p≤ 0.05 were considered significant.

### Reagents for in vitro assay

**Reagents for in vitro assay:**
| **<u>Treatment</u>** | **<u>Working Concentration</u>** | **<u>Catalogue #</u>** |
| --- | --- | --- |
| IL-4 | 30 ng/µL | R&D; 404ML010 |
| IL-13 | 30 ng/µL | R&D; 413ML025 |
| M-CSF | 20 ng/µL | R&D; 416ML010 |
| Paraffin wax |  | SIGMA |
| Anti-DCSTAMP blocking Ab | 20 µg/µL | Millipore Sigma; MABF39-1 (Clone 1A2) |
| Anti-CD81 blocking Ab | 50 µg/µL | Genetex; GTX101766 |
| Cytochalasin D | 1µM | Tocris Bioscience |
| Cycloheximide | 100 µg/mL | Millipore Sigma |
| MG132 | 5-10 µM | Tocris Bioscience |

**Immunofluorescent reagents:**
| <u>Antibody/stain</u> | <u>Working Concentration</u> | <u>Catalogue #</u> |
| --- | --- | --- |
| Mac-3 | 1:200 | BD Pharmingen; 550292 |
| CD13, clone SL13 | 1:500 | Millipore Sigma; MABC950 |
| CD9 | 1:200 | MyBiosource; MBS857046 |
| Phalloidin | 1:1000 | Invitrogen; R415 |
| CUL4 | 1:200 | Kerafast; EB0022-FP |
| DCSTAMP | 1:100 | Millipore; MABF39-I |
| DAPI | 1:1000 | Invitrogen; D1306 |
| Alexa Fluor Secondary Ab 488 and 594 | 1:1000 | Invitrogen |

**Flow Cytometry reagents:**
| <u>Antibody/stain</u> | <u>Working Concentration</u> | <u>Catalog #</u> |
| --- | --- | --- |
| anti-CD3 APC | 1:20 | Biolegend; 100235 |
| anti-B220 APC | 1:80 | Biolegend; 103212 |
| anti-NK1.1 APC | 1:20 | Biolegend; 108710 |
| anti-CD11b 450 | 1:20 | Invitrogen; 48-0112-82 |
| anti-CD115 PE/Cy7 | 1:80 | Biolegend; 135524 |
| anti-Ly6C AF700 | 1:200 | Biolegend; 128024 |

## Discussion

Implantable medical devices have become invaluable tools to prevent, monitor, treat, diagnose or alleviate numerous health conditions and serve to improve our overall quality of life [1]. Not surprisingly, these devices comprise a large and rapidly growing sector of the health care market and considering our aging population, their demand will only increase. Implantable devices span all medical specialties and include artificial joints, heart valves, cardiac pacemakers, biologic meshes, cardiac and glucose sensors, insulin infusion pumps, breast, dental and cochlear *implants* and intraocular lenses among hundreds of others. As these implants invariably invoke a deleterious response from the host, it is imperative to focus on the FBR as a fundamental barrier that must be surmounted to allow critical advancements and improvements to life-enhancing medical devices (*83*). Therefore, uncovering the fundamental mechanisms regulating the FBR will identify the novel molecules and pathways that represent tomorrow’s therapeutic targets.

Monocytes and macrophages have many functions ranging from phagocytosis of invading pathogens, triggering the adaptive immune response, to regulating lipid and iron metabolism (84). Certain subsets of macrophages are able to fuse their membranes, forming syncytial multinucleated giant cells (MGCs) that share a common cytoplasm and plasma membrane (85). The three primary MGC subsets are osteoclasts, foreign body giant cells (FBGCs), and Langhans giant cells (86), which represent distinct lineages and result from distinct signaling pathways. MGC fusion with tumor cells. Osteoclast fusion is primarily regulated by the receptor activator of nuclear factor-kappa-B ligand (RANKL) signaling pathway. RANKL is produced by osteoblasts and stromal cells and binds to its receptor RANK on preosteoclasts, promoting their fusion into multinucleated osteoclasts. Osteoclast fusion is a physiological process that enables bone remodeling by breaking down and resorbing bone tissue. In contrast, foreign body-induced fusion is a pathological process triggered by the presence of foreign materials that activate macrophages and monocytes to fuse and form FBGCs which phagocytose and remove foreign substances. Formation of both of these lineages can be recapitulated *in vitro* by stimulation of bone marrow progenitors with appropriate cytokines, RANKL for osteoclasts and IL-4 and IL-13 for FBGC-like MGCs (42). While osteoclast fusion has been intensely studied, little is known about fusion in other MGC subtypes (14). However, since the basis for revision surgery in nearly 55% of hip (87) and 60.2% of knee revisions (88) is aseptic loosening, it is imperative to optimize primary outcomes in implantable devices by controlling the FBR.

The mechanism of the FBR depends on the physical and biochemical properties of the implant, the implant site and host immune responses (48). It is a complex multi-factorial process, and to date, which components of the FBR are causal to and/or predictive of implant failure remain unresolved (8, 9, 89). Despite the fact that macrophages are key players in inflammation, studies focusing on preventing FBR at the stage of inflammation mediated by FBGCs are lacking, partly due to some ambiguity as to the true contribution of FBGCs and the controversies surrounding their role in FBR. Numerous investigations have indicated that FBGCs contribute to the degradation of foreign particles through secretion of ROS, MMPs, and acids (10), and are a source of cytokines and chemokines (11, 12), and ROS (13, 14). Alternatively, two studies found that FBGCs were dispensable for FBR-induced encapsulation (8, 9) where both CCL-2 and MMP-9 null mice had reduced FBGC numbers but normal capsule thickness, although MMP-9 null mice reportedly showed defects in collagen deposition and assembly, illustrating the complexity of the FBR. The increased number of FBGCs at the peri-implant site in our CD13^KO^ model is accompanied by increased classic fibrosis and amplified intracellular and systemic TGF-β levels, indicative of a more intense FBR. This would imply that FBGCs are a critical, causal component of FBR progression *in vivo* and supports CD13 as a potential target in minimizing the deleterious effects of the FBR. Alternatively, it is possible that a lack of CD13 could alter the FBGC phenotype, perhaps making it more aggressive, which also lead to exacerbated fibrosis independent of FBGC numbers. Further analysis to uncover potential differences in the transcriptional profiles of the cell types at the implant site in WT vs, CD13^KO^ mice by sc-RNA seq is currently underway in our laboratory to address this possibility.

Cell-cell fusion is tightly controlled at the molecular level, involving the coordination of multiple factors, some that are shared and others specific to the cell and tissue type. While various cell types and biological events involve cell fusion, the process has yet to be reduced to a well-ordered scheme. However, all fusion requires the initial steps of cell migration toward neighboring cells; organization, recognition and engagement of membrane adhesion molecules (90); and the assembly of thin, actin-based protrusions extending between cells that colocalize with actin-polymerization proteins (91). These steps also depend on the spatial and proximate coordination of components of key signaling pathways and their target adhesion molecules, cell surface receptors, intracellular kinases and transcription factors, together culminating in the fusion event (92). We have shown that CD13 is a cell-cell adhesion molecule (33–35, 93); regulates cell migration (38, 59), controls the assembly of actin structures (38, 94) and directs osteoclast fusion (39) and thus could contribute to FBGC fusion at a number of steps. However, we have also demonstrated that in some of these cases, CD13 acts as a transmembrane anchor to bind and properly position signaling complexes in the correct membrane context to enable subsequent steps. Anchor proteins in turn often bind to scaffold proteins that assemble various signaling components into functional signaling complexes. We have shown that CD13 promoted ARF6 GTPase activity by binding to the scaffold IQGAP and positioning the IQGAP/ARF6/EFA6 signaling complex at the leading edge of the cell membrane, promoting ARF6 GTPase cycling and cell migration (38). In this regard, we consider that CD13 as a transmembrane anchor that coordinates various signaling steps that allow fusion to occur. This is supported by the clear mislocalization of the CUL4, DCSTAMP1 and CD9 proteins in CD13^KO^ FBGCs, suggesting that CD13-dependent membrane tethering of the CUL4 complex is required for fusogen degradation and normal cell-cell fusion.

## Supporting information

Manuscript file with figures

## Author contributions

MG and LHS designed the research plan. MG, FM, RKV, RN, EJ and SN conducted experiments and MG, FM, RN, AD, SN, EJ and PM analyzed data. MG and LHS wrote the manuscript.

## Acknowledgments

We thank Ms. Keriahen Morais for technical assistance. We also thank Dr. Kevin Claffey for his advice and help with implant harvest and use of the histology core at UConn Health. We thank Dr. Zhifang Hao for the FPPE sections of mesh implants and trichrome staining. This work was supported by National Institutes of Health grant 1R21AI15 (LHS and MG) and Research Excellence grant, UCONN (LHS and MG).

---

**FIG S1. Step-wise flow-sorting of myeloid progenitors from WT and CD13^KO^ BM. A.** Live (CD3/B220/Nk1.1)^-^ CD11b^lo/-^ CD115^hi^ Ly6G^+^ BM progenitor cells were sorted to homogeneity and analyzed by FACS Diva. **B**. Giemsa analysis of flow sorted BM progenitor cells differentiated in presence of MCSF + IL-4 + IL-13 for d5 showed equivalent number of total nuclei/field between genotypes. Data represents +/- SD of 3 independent experiments. N=3/genotype.

**FIG S2. A-B. CD13 is highly expressed in WT MGC but not in CD13^KO^ counterpart.** CD13 (SL13 mAb; red) expression in the WT membrane in (**A**) multinucleated giant cells at d5 post fusion compared to CD13^KO^ (**B**) cells under similar condition. **C-D**. Secondary Ab controls in WT cells undergoing fusion.

**FIG S3. Actin-based protrusion prefusion drives hyperfusion phenotype in CD13^KO^. A.** Average length of protrusion of giemsa stained flow sorted BM derived myeloid cells stimulated with MCSF + IL-4 + IL-13. **B.** IF analysis of Phalloidin stained thioglycollate elicited WT and CD13^KO^ peritoneal macrophages grown on treated optical glass for d2 post MCSF+ IL-4+ IL-13 treatment exhibited enhanced actin-based protrusions in CD13^KO^ compared to WT cells. **C**. % Fusion index is significantly reduced in both WT and CD13^KO^ TG-macs treated with Cytochalasin D (CytoD; 1μM) at d1 under fusogenic condition compared to vehicle control. Data represents +/- SD of 3 independent experiments. N=3/genotype. **;p<0.01, *; p<0.05. Magnification; 63X oil. Scale bar; 10µM.

**FIG S4. Expression of fusion promoting transcripts are equivalent among genotypes.** WT and CD13^KO^ flow sorted BM progenitor cells were stimulated with MCSF + IL-4 + IL-13 over time (d0, d3 and d5) and expression of fusion regulatory transcripts were analyzed by quantitative RT-PCR. Relative transcript level normalized to GAPDH indicated equivalent levels of *DCSTAMP*, *CD9* and CD81 genes in both genotypes. Data represents +/- SD of 3 independent experiments. N=3/genotype.

**FIG S5. Post-transcriptional regulation of fusion promoting proteins by CD13 mediates hyperfusion phenotype in absence of CD13.** WT and CD13^KO^ BM-derived myeloid progenitors under fusogenic conditions were treated with cycloheximide, (CHX; protein synthesis inhibitor) (100μg/ml) +/- 5 & 10μM MG132, (proteasomal degradation inhibitor) for 8h and surface expression analyzed by flow cytometry. Representative histogram depicts surface mean fluorescence intensity of CD9 (**A**) and DCSTAMP (**B**) in WT and CD13^KO^ in presence of CHX vs. CHX + MG132. MG132-mediated blocking of proteasomal inhibition led to significantly delayed surface expression of fusogens in WT compared to CD13^KO^ cells +/- CHX treatment. Data represents +/- SD of 2 independent experiments. N=3/genotype.

**FIG S6. Cullin 4A and DCSTAMP are mislocalized in the absence of CD13.** IF analysis (**A**), zoomed image (**B**) and Pearson’s correlation coefficient (**C**) of WT and CD13^KO^ TG-peritoneal macrophages under fusogenic condition at d3 indicated that CUL-4A (red) and DCSTAMP (green) do not colocalize in CD13^KO^ cells compared to WT cells. Data represents +/- SD of 2 independent experiments.

**FIG S7. Anpep (CD13) is one of the most highly induced genes in FBGCs compared to immediate precursors by transcriptome analysis of single cell RNA seq**. Violin plots of single cell RNA seq in implant induced FBR indicated that the distribution and expression levels of Anpep (CD13), is highly induced in FBGCs (multinucleated giant cells) compared to its immediate precursors (IP) and M2 macrophages (M2).

## References

1. Skokos, E. A., A. Charokopos, K. Khan, J. Wanjala, and T. R. Kyriakides. 2011. Lack of TNF-alpha-induced MMP-9 production and abnormal E-cadherin redistribution associated with compromised fusion in MCP-1-null macrophages. Am J Pathol 178: 2311–2321.

2. Petrany, M. J., and D. P. Millay. 2019. Cell Fusion: Merging Membranes and Making Muscle. Trends Cell Biol 29: 964–973.

3. Brukman, N. G., B. Uygur, B. Podbilewicz, and L. V. Chernomordik. 2019. How cells fuse. J Cell Biol 218: 1436–1451.

4. McNally, A. K., and J. M. Anderson. 2005. Multinucleated giant cell formation exhibits features of phagocytosis with participation of the endoplasmic reticulum. Exp Mol Pathol 79: 126–135.

5. Jay, S. M., E. Skokos, F. Laiwalla, M. M. Krady, and T. R. Kyriakides. 2007. Foreign body giant cell formation is preceded by lamellipodia formation and can be attenuated by inhibition of Rac1 activation. Am J Pathol 171: 632–640.

6. Anderson, J. M., A. Rodriguez, and D. T. Chang. 2008. Foreign body reaction to biomaterials. Semin Immunol 20: 86–100.

7. Sheikh, Z., P. J. Brooks, O. Barzilay, N. Fine, and M. Glogauer. 2015. Macrophages, Foreign Body Giant Cells and Their Response to Implantable Biomaterials. Materials (Basel) 8: 5671–5701.

8. Kyriakides, T. R., M. J. Foster, G. E. Keeney, A. Tsai, C. M. Giachelli, I. Clark-Lewis, B. J. Rollins, and P. Bornstein. 2004. The CC chemokine ligand, CCL2/MCP1, participates in macrophage fusion and foreign body giant cell formation. Am J Pathol 165: 2157–2166.

9. MacLauchlan, S., E. A. Skokos, N. Meznarich, D. H. Zhu, S. Raoof, J. M. Shipley, R. M. Senior, P. Bornstein, and T. R. Kyriakides. 2009. Macrophage fusion, giant cell formation, and the foreign body response require matrix metalloproteinase 9. J Leukoc Biol 85: 617–626.

10. Jones, J. A., D. T. Chang, H. Meyerson, E. Colton, I. K. Kwon, T. Matsuda, and J. M. Anderson. 2007. Proteomic analysis and quantification of cytokines and chemokines from biomaterial surface-adherent macrophages and foreign body giant cells. J Biomed Mater Res A 83: 585–596.

11. Hernandez-Pando, R., Q. L. Bornstein, D. Aguilar Leon, E. H. Orozco, V. K. Madrigal, and E. Martinez Cordero. 2000. Inflammatory cytokine production by immunological and foreign body multinucleated giant cells. Immunology 100: 352–358.

12. Saleh, L. S., and S. J. Bryant. 2017. In Vitro and In Vivo Models for Assessing the Host Response to Biomaterials. Drug Discov Today Dis Models 24: 13–21.

13. Enelow, R. I., G. W. Sullivan, H. T. Carper, and G. L. Mandell. 1992. Cytokine-induced human multinucleated giant cells have enhanced candidacidal activity and oxidative capacity compared with macrophages. J Infect Dis 166: 664–668.

14. Ahmadzadeh, K., M. Vanoppen, C. D. Rose, P. Matthys, and C. H. Wouters. 2022. Multinucleated Giant Cells: Current Insights in Phenotype, Biological Activities, and Mechanism of Formation. Front Cell Dev Biol 10: 873226.

15. Lee, J. H., C. F. Hsieh, H. W. Liu, C. Y. Chen, S. C. Wu, T. W. Chen, C. S. Hsu, Y. H. Liao, C. Y. Yang, J. F. Shyu, W. B. Fischer, and C. H. Lin. 2017. Lipid raft-associated stomatin enhances cell fusion. FASEB J 31: 47–59.

16. Ishii, M., K. Iwai, M. Koike, S. Ohshima, E. Kudo-Tanaka, T. Ishii, T. Mima, Y. Katada, K. Miyatake, Y. Uchiyama, and Y. Saeki. 2006. RANKL-induced expression of tetraspanin CD9 in lipid raft membrane microdomain is essential for cell fusion during osteoclastogenesis. J Bone Miner Res 21: 965–976.

17. Ha, H., H. B. Kwak, S. K. Lee, D. S. Na, C. E. Rudd, Z. H. Lee, and H. H. Kim. 2003. Membrane rafts play a crucial role in receptor activator of nuclear factor kappaB signaling and osteoclast function. J Biol Chem 278: 18573–18580.

18. Hogue, I. B., J. R. Grover, F. Soheilian, K. Nagashima, and A. Ono. 2011. Gag induces the coalescence of clustered lipid rafts and tetraspanin-enriched microdomains at HIV-1 assembly sites on the plasma membrane. J Virol 85: 9749–9766.

19. Shin, N. Y., H. Choi, L. Neff, Y. Wu, H. Saito, S. M. Ferguson, P. De Camilli, and R. Baron. 2014. Dynamin and endocytosis are required for the fusion of osteoclasts and myoblasts. J Cell Biol 207: 73–89.

20. Yagi, M., T. Miyamoto, Y. Toyama, and T. Suda. 2006. Role of DC-STAMP in cellular fusion of osteoclasts and macrophage giant cells. J Bone Miner Metab 24: 355–358.

21. Kukita, T., N. Wada, A. Kukita, T. Kakimoto, F. Sandra, K. Toh, K. Nagata, T. Iijima, M. Horiuchi, H. Matsusaki, K. Hieshima, O. Yoshie, and H. Nomiyama. 2004. RANKL-induced DC-STAMP is essential for osteoclastogenesis. J Exp Med 200: 941–946.

22. Song, R. L., X. Z. Liu, J. Q. Zhu, J. M. Zhang, Q. Gao, H. Y. Zhao, A. Z. Sheng, Y. Yuan, J. H. Gu, H. Zou, Q. C. Wang, and Z. P. Liu. 2014. New roles of filopodia and podosomes in the differentiation and fusion process of osteoclasts. Genet Mol Res 13: 4776–4787.

23. Yagi, M., T. Miyamoto, Y. Sawatani, K. Iwamoto, N. Hosogane, N. Fujita, K. Morita, K. Ninomiya, T. Suzuki, K. Miyamoto, Y. Oike, M. Takeya, Y. Toyama, and T. Suda. 2005. DC-STAMP is essential for cell-cell fusion in osteoclasts and foreign body giant cells. J Exp Med 202: 345–351.

24. Miyamoto, H., T. Suzuki, Y. Miyauchi, R. Iwasaki, T. Kobayashi, Y. Sato, K. Miyamoto, H. Hoshi, K. Hashimoto, S. Yoshida, W. Hao, T. Mori, H. Kanagawa, E. Katsuyama, A. Fujie, H. Morioka, M. Matsumoto, K. Chiba, M. Takeya, Y. Toyama, and T. Miyamoto. 2012. Osteoclast stimulatory transmembrane protein and dendritic cell-specific transmembrane protein cooperatively modulate cell-cell fusion to form osteoclasts and foreign body giant cells. J Bone Miner Res 27: 1289–1297.

25. Ohnami, N., A. Nakamura, M. Miyado, M. Sato, N. Kawano, K. Yoshida, Y. Harada, Y. Takezawa, S. Kanai, C. Ono, Y. Takahashi, K. Kimura, T. Shida, K. Miyado, and A. Umezawa. 2012. CD81 and CD9 work independently as extracellular components upon fusion of sperm and oocyte. Biol Open 1: 640–647.

26. Champion, T. C., L. J. Partridge, S. M. Ong, B. Malleret, S. C. Wong, and P. N. Monk. 2018. Monocyte Subsets Have Distinct Patterns of Tetraspanin Expression and Different Capacities to Form Multinucleate Giant Cells. Front Immunol 9: 1247.

27. Singethan, K., and J. Schneider-Schaulies. 2008. Tetraspanins: Small transmembrane proteins with big impact on membrane microdomain structures. Communicative & integrative biology 1: 11–13.

28. Kim, K., S. H. Lee, J. Ha Kim, Y. Choi, and N. Kim. 2008. NFATc1 induces osteoclast fusion via up-regulation of Atp6v0d2 and the dendritic cell-specific transmembrane protein (DC-STAMP). Mol Endocrinol 22: 176–185.

29. Moller, A. M., J. M. Delaisse, and K. Soe. 2017. Osteoclast Fusion: Time-Lapse Reveals Involvement of CD47 and Syncytin-1 at Different Stages of Nuclearity. J Cell Physiol 232: 1396–1403.

30. Li, F., H. Chung, S. V. Reddy, G. Lu, N. Kurihara, A. Z. Zhao, and G. D. Roodman. 2005. Annexin II stimulates RANKL expression through MAPK. J Bone Miner Res 20: 1161–1167.

31. Erlandsson, M. C., M. D. Svensson, I. M. Jonsson, L. Bian, N. Ambartsumian, S. Andersson, Z. Peng, J. Vaaraniemi, C. Ohlsson, K. M. E. Andersson, and M. I. Bokarewa. 2013. Expression of metastasin S100A4 is essential for bone resorption and regulates osteoclast function. Biochim Biophys Acta 1833: 2653–2663.

32. Verma, S. K., E. Leikina, K. Melikov, C. Gebert, V. Kram, M. F. Young, B. Uygur, and L. V. Chernomordik. 2018. Cell-surface phosphatidylserine regulates osteoclast precursor fusion. J Biol Chem 293: 254–270.

33. Mina-Osorio, P., B. Winnicka, C. O’Conor, C. L. Grant, L. K. Vogel, D. Rodriguez-Pinto, K. V. Holmes, E. Ortega, and L. H. Shapiro. 2008. CD13 is a novel mediator of monocytic/endothelial cell adhesion. J Leukoc Biol 84: 448–459.

34. Ghosh, M., C. Gerber, M. M. Rahman, K. M. Vernier, F. E. Pereira, J. Subramani, L. A. Caromile, and L. H. Shapiro. 2014. Molecular mechanisms regulating CD13-mediated adhesion. Immunology 142: 636–647.

35. Subramani, J., M. Ghosh, M. M. Rahman, L. A. Caromile, C. Gerber, K. Rezaul, D. K. Han, and L. H. Shapiro. 2013. Tyrosine Phosphorylation of CD13 Regulates Inflammatory Cell-Cell Adhesion and Monocyte Trafficking. J Immunol 191: 3905–3912.

36. Rahman, M. M., M. Ghosh, J. Subramani, G. H. Fong, M. E. Carlson, and L. H. Shapiro. 2014. CD13 regulates anchorage and differentiation of the skeletal muscle satellite stem cell population in ischemic injury. Stem Cells 32: 1564–1577.

37. Ghosh, M., B. McAuliffe, J. Subramani, S. Basu, and L. H. Shapiro. 2012. CD13 Regulates Dendritic Cell Cross-Presentation and T Cell Responses by Inhibiting Receptor-Mediated Antigen Uptake. J Immunol 188: 5489–5499.

38. Ghosh, M., R. Lo, I. Ivic, B. Aguilera, V. Qendro, C. Devarakonda, and L. H. Shapiro. 2019. CD13 tethers the IQGAP1-ARF6-EFA6 complex to the plasma membrane to promote ARF6 activation, beta1 integrin recycling, and cell migration. Sci Signal 12.

39. Ghosh, M., T. Kelava, I. V. Madunic, I. Kalajzic, and L. H. Shapiro. 2021. CD13 is a critical regulator of cell-cell fusion in osteoclastogenesis. Sci Rep 11: 10736.

40. Zhang, Y., G. Morrone, J. Zhang, X. Chen, X. Lu, L. Ma, M. Moore, and P. Zhou. 2003. CUL-4A stimulates ubiquitylation and degradation of the HOXA9 homeodomain protein. EMBO J 22: 6057–6067.

41. Bulatov, E., and A. Ciulli. 2015. Targeting Cullin-RING E3 ubiquitin ligases for drug discovery: structure, assembly and small-molecule modulation. Biochem J 467: 365–386.

42. Kloc, M., A. Subuddhi, A. Uosef, J. Z. Kubiak, and R. M. Ghobrial. 2022. Monocyte-Macrophage Lineage Cell Fusion. Int J Mol Sci 23.

43. Faust, J. J., W. Christenson, K. Doudrick, R. Ros, and T. P. Ugarova. 2017. Development of fusogenic glass surfaces that impart spatiotemporal control over macrophage fusion: Direct visualization of multinucleated giant cell formation. Biomaterials 128: 160–171.

44. Enelow, R. I., G. W. Sullivan, H. T. Carper, and G. L. Mandell. 1992. Induction of multinucleated giant cell formation from in vitro culture of human monocytes with interleukin-3 and interferon-gamma: comparison with other stimulating factors. Am J Respir Cell Mol Biol 6: 57–62.

45. Klopfleisch, R., and F. Jung. 2017. The pathology of the foreign body reaction against biomaterials. J Biomed Mater Res A 105: 927–940.

46. Carnicer-Lombarte, A., S. T. Chen, G. G. Malliaras, and D. G. Barone. 2021. Foreign Body Reaction to Implanted Biomaterials and Its Impact in Nerve Neuroprosthetics. Front Bioeng Biotechnol 9: 622524.

47. Novitsky, Y. W., S. B. Orenstein, and D. L. Kreutzer. 2014. Comparative analysis of histopathologic responses to implanted porcine biologic meshes. Hernia 18: 713–721.

48. Ibrahim, M., J. Bond, M. A. Medina, L. Chen, C. Quiles, G. Kokosis, L. Bashirov, B. Klitzman, and H. Levinson. 2017. Characterization of the Foreign Body Response to Common Surgical Biomaterials in a Murine Model. Eur J Plast Surg 40: 383–392.

49. Wang, Y., P. J. Brooks, J. J. Jang, A. S. Silver, P. D. Arora, C. A. McCulloch, and M. Glogauer. 2015. Role of actin filaments in fusopod formation and osteoclastogenesis. Biochim Biophys Acta 1853: 1715–1724.

50. Shilagardi, K., S. Li, F. Luo, F. Marikar, R. Duan, P. Jin, J. H. Kim, K. Murnen, and E. H. Chen. 2013. Actin-propelled invasive membrane protrusions promote fusogenic protein engagement during cell-cell fusion. Science 340: 359–363.

51. Kim, Y. S., S. Shin, E. J. Choi, S. W. Moon, C. K. Jung, Y. J. Chung, and S. H. Lee. 2022. Different Molecular Features of Epithelioid and Giant Cells in Foreign Body Reaction Identified by Single-Cell RNA Sequencing. J Invest Dermatol 142: 3232–3242 e3216.

52. Lignitto, L., and M. Pagano. 2021. Linking ubiquitin to actin dynamics during cell fusion. Dev Cell 56: 569–570.

53. Potgens, A. J., S. Drewlo, M. Kokozidou, and P. Kaufmann. 2004. Syncytin: the major regulator of trophoblast fusion? Recent developments and hypotheses on its action. Hum Reprod Update 10: 487–496.

54. Chen, E. H., and E. N. Olson. 2005. Unveiling the mechanisms of cell-cell fusion. Science 308: 369–373.

55. Dugast, C., A. Gaudin, and L. Toujas. 1997. Generation of multinucleated giant cells by culture of monocyte-derived macrophages with IL-4. Jr.of Leukocyte Biology 61: 517–521.

56. Oikawa, T., and K. Matsuo. 2012. Possible role of IRTKS in Tks5-driven osteoclast fusion. Communicative & integrative biology 5: 511–515.

57. DeFife, K. M., C. R. Jenney, E. Colton, and J. M. Anderson. 1999. Disruption of filamentous actin inhibits human macrophage fusion. FASEB J 13: 823–832.

58. Licona-Limon, I., C. A. Garay-Canales, O. Munoz-Paleta, and E. Ortega. 2015. CD13 mediates phagocytosis in human monocytic cells. J Leukoc Biol 98: 85–98.

59. Petrovic, N., W. Schacke, J. R. Gahagan, C. A. O’Conor, B. Winnicka, R. E. Conway, P. Mina-Osorio, and L. H. Shapiro. 2007. CD13/APN regulates endothelial invasion and filopodia formation. Blood 110: 142–150.

60. Norris, R. P., V. Baena, and M. Terasaki. 2017. Localization of phosphorylated connexin 43 using serial section immunogold electron microscopy. J Cell Sci 130: 1333–1340.

61. Norris, R. P., and M. Terasaki. 2021. Gap junction internalization and processing in vivo: a 3D immuno-electron microscopy study. J Cell Sci 134.

62. Jimenez, N., E. G. Van Donselaar, D. A. De Winter, K. Vocking, A. J. Verkleij, and J. A. Post. 2010. Gridded Aclar: preparation methods and use for correlative light and electron microscopy of cell monolayers, by TEM and FIB-SEM. J Microsc 237: 208–220.

63. Takeda, Y., I. Tachibana, K. Miyado, M. Kobayashi, T. Miyazaki, T. Funakoshi, H. Kimura, H. Yamane, Y. Saito, H. Goto, T. Yoneda, M. Yoshida, T. Kumagai, T. Osaki, S. Hayashi, I. Kawase, and E. Mekada. 2003. Tetraspanins CD9 and CD81 function to prevent the fusion of mononuclear phagocytes. J Cell Biol 161: 945–956.

64. Parthasarathy, V., F. Martin, A. Higginbottom, H. Murray, G. W. Moseley, R. C. Read, G. Mal, R. Hulme, P. N. Monk, and L. J. Partridge. 2009. Distinct roles for tetraspanins CD9, CD63 and CD81 in the formation of multinucleated giant cells. Immunology 127: 237–248.

65. Fanaei, M., P. N. Monk, and L. J. Partridge. 2011. The role of tetraspanins in fusion. Biochem Soc Trans 39: 524–528.

66. Warren, G. L., T. Hulderman, D. Mishra, X. Gao, L. Millecchia, L. O’Farrell, W. A. Kuziel, and P. P. Simeonova. 2005. Chemokine receptor CCR2 involvement in skeletal muscle regeneration. FASEB J 19: 413–415.

67. Chiu, Y. H., K. A. Mensah, E. M. Schwarz, Y. Ju, M. Takahata, C. Feng, L. A. McMahon, D. G. Hicks, B. Panepento, P. C. Keng, and C. T. Ritchlin. 2012. Regulation of human osteoclast development by dendritic cell-specific transmembrane protein (DC-STAMP). J Bone Miner Res 27: 79–92.

68. Rusilowicz-Jones, E. V., S. Urbe, and M. J. Clague. 2022. Protein degradation on the global scale. Mol Cell 82: 1414–1423.

69. Chang, Y., and S. C. Finnemann. 2007. Tetraspanin CD81 is required for the alpha v beta5-integrin-dependent particle-binding step of RPE phagocytosis. J Cell Sci 120: 3053–3063.

70. Mensah, K. A., C. T. Ritchlin, and E. M. Schwarz. 2010. RANKL induces heterogeneous DC-STAMP(lo) and DC-STAMP(hi) osteoclast precursors of which the DC-STAMP(lo) precursors are the master fusogens. J Cell Physiol 223: 76–83.

71. Rodriguez-Perez, F., A. G. Manford, A. Pogson, A. J. Ingersoll, B. Martinez-Gonzalez, and M. Rape. 2021. Ubiquitin-dependent remodeling of the actin cytoskeleton drives cell fusion. Dev Cell 56: 588–601 e589.

72. Kanemoto, S., Y. Kobayashi, T. Yamashita, T. Miyamoto, M. Cui, R. Asada, X. Cui, K. Hino, M. Kaneko, T. Takai, K. Matsuhisa, N. Takahashi, and K. Imaizumi. 2015. Luman is involved in osteoclastogenesis through the regulation of DC-STAMP expression, stability and localization. J Cell Sci 128: 4353–4365.

73. Clague, M. J., and S. Urbe. 2010. Ubiquitin: same molecule, different degradation pathways. Cell 143: 682–685.

74. Cockram, P. E., M. Kist, S. Prakash, S. H. Chen, I. E. Wertz, and D. Vucic. 2021. Ubiquitination in the regulation of inflammatory cell death and cancer. Cell Death Differ 28: 591–605.

75. Damgaard, R. B. 2021. The ubiquitin system: from cell signalling to disease biology and new therapeutic opportunities. Cell Death Differ 28: 423–426.

76. Zheng, G., J. Zhang, H. Zhao, H. Wang, M. Pang, X. Qiao, S. R. Lee, T. T. Hsu, T. K. Tan, J. G. Lyons, Y. Zhao, X. Tian, D. A. F. Loebel, I. Rubera, M. Tauc, Y. Wang, Y. Wang, Y. M. Wang, Q. Cao, C. Wang, V. W. S. Lee, S. I. Alexander, P. P. L. Tam, and D. C. H. Harris. 2016. alpha3 Integrin of Cell-Cell Contact Mediates Kidney Fibrosis by Integrin-Linked Kinase in Proximal Tubular E-Cadherin Deficient Mice. Am J Pathol 186: 1847–1860.

77. Sun, T., Z. Liu, and Q. Yang. 2020. The role of ubiquitination and deubiquitination in cancer metabolism. Mol Cancer 19: 146.

78. Sarikas, A., T. Hartmann, and Z. Q. Pan. 2011. The cullin protein family. Genome biology 12: 220.

79. Chen, Z., W. Zhang, K. Jiang, B. Chen, K. Wang, L. Lao, C. Hou, F. Wang, C. Zhang, and H. Shen. 2018. MicroRNA-300 Regulates the Ubiquitination of PTEN through the CRL4B(DCAF13) E3 Ligase in Osteosarcoma Cells. Mol Ther Nucleic Acids 10: 254–268.

80. Xin, M., X. Jin, X. Cui, C. Jin, L. Piao, Y. Wan, S. Xu, S. Zhang, X. Yue, H. Wang, Y. Nan, and X. Cheng. 2019. Dipeptidyl peptidase-4 inhibition prevents vascular aging in mice under chronic stress: Modulation of oxidative stress and inflammation. Chemico-biological interactions 314: 108842.

81. Sweeney, M. A., P. Iakova, L. Maneix, F. Y. Shih, H. E. Cho, E. Sahin, and A. Catic. 2020. The ubiquitin ligase Cullin-1 associates with chromatin and regulates transcription of specific c-MYC target genes. Sci Rep 10: 13942.

82. Ghosh, M., R. Lo, I. Ivic, B. Aguilera, V. Qendro, C. Devarakonda, and L. H. Shapiro. 2019. CD13 Targets IQGAP1-ARF6-EFA6 to the Plasma Membrane to Promote ARF6 Activation, Coordinate β1-Integrin Recycling and Cell Migration. Science Signaling revision under review.

83. Didyuk, O., N. Econom, A. Guardia, K. Livingston, and U. Klueh. 2021. Continuous Glucose Monitoring Devices: Past, Present, and Future Focus on the History and Evolution of Technological Innovation. J Diabetes Sci Technol 15: 676–683.

84. Lazarov, T., S. Juarez-Carreno, N. Cox, and F. Geissmann. 2023. Physiology and diseases of tissue-resident macrophages. Nature 618: 698–707.

85. Pereira, M., E. Petretto, S. Gordon, J. H. D. Bassett, G. R. Williams, and J. Behmoaras. 2018. Common signalling pathways in macrophage and osteoclast multinucleation. J Cell Sci 131.

86. Liu, L., W. F. Anderson, R. W. Beart, E. M. Gordon, and F. L. Hall. 2000. Incorporation of tumor vasculature targeting motifs into moloney murine leukemia virus env escort proteins enhances retrovirus binding and transduction of human endothelial cells. J Virol 74: 5320–5328.

87. Cherian, J. J., J. J. Jauregui, S. Banerjee, T. Pierce, and M. A. Mont. 2015. What Host Factors Affect Aseptic Loosening After THA and TKA? Clin Orthop Relat Res 473: 2700–2709.

88. Baek, J. H., S. C. Lee, S. Ryu, H. S. Ahn, and C. H. Nam. 2022. Early aseptic loosening of primary total knee arthroplasty in patients with osteonecrosis of the knee: A case series. Clin Case Rep 10: e6773.

89. Chen, Y., K. J. Pandya, O. Hyrien, P. C. Keng, T. Smudzin, J. Anderson, R. Qazi, B. Smith, T. J. Watson, R. H. Feins, and D. W. Johnstone. 2011. Preclinical and pilot clinical studies of docetaxel chemoradiation for Stage III non-small-cell lung cancer. Int J Radiat Oncol Biol Phys 80: 1358–1364.

90. McNally, A. K., and J. M. Anderson. 2002. Beta1 and beta2 integrins mediate adhesion during macrophage fusion and multinucleated foreign body giant cell formation. Am J Pathol 160: 621–630.

91. Zito, F., N. Lampiasi, I. Kireev, and R. Russo. 2016. United we stand: Adhesion and molecular mechanisms driving cell fusion across species. Eur J Cell Biol 95: 552–562.

92. Zhou, X., and J. L. Platt. 2011. Molecular and cellular mechanisms of mammalian cell fusion. Adv Exp Med Biol 713: 33–64.

93. Mina-Osorio, P., L. H. Shapiro, and E. Ortega. 2006. CD13 in cell adhesion: aminopeptidase N (CD13) mediates homotypic aggregation of monocytic cells. J Leukoc Biol 79: 719–730.

94. Ghosh, M., J. Subramani, M. M. Rahman, and L. H. Shapiro. 2015. CD13 Restricts TLR4 Endocytic Signal Transduction in Inflammation. J Immunol 194: 4466–4476.

