## Supplementary figures and images for "The Implant-Induced Foreign Body Response is Limited by CD13-Dependent Regulation of Ubiquitination of Fusogenic Proteins"

### Manuscript file with figures

FIG S1

**A**

**Live (CD3/B220/Nk1.1)[-] CD11b[lo/-] CD115[hi] Ly6G[+] BM progenitor cells**

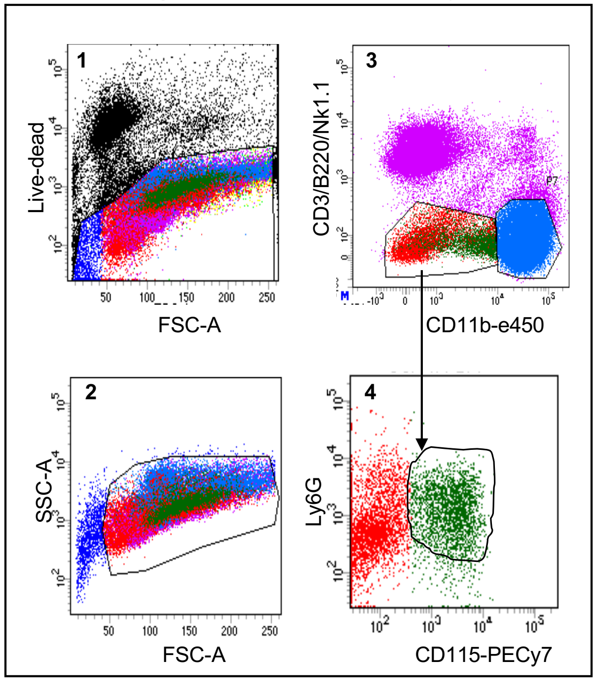


**B**

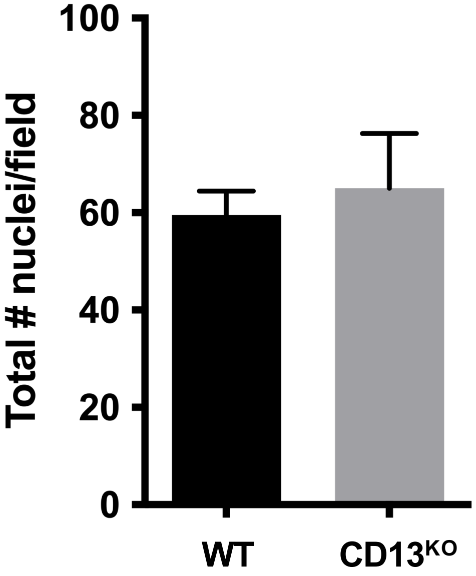

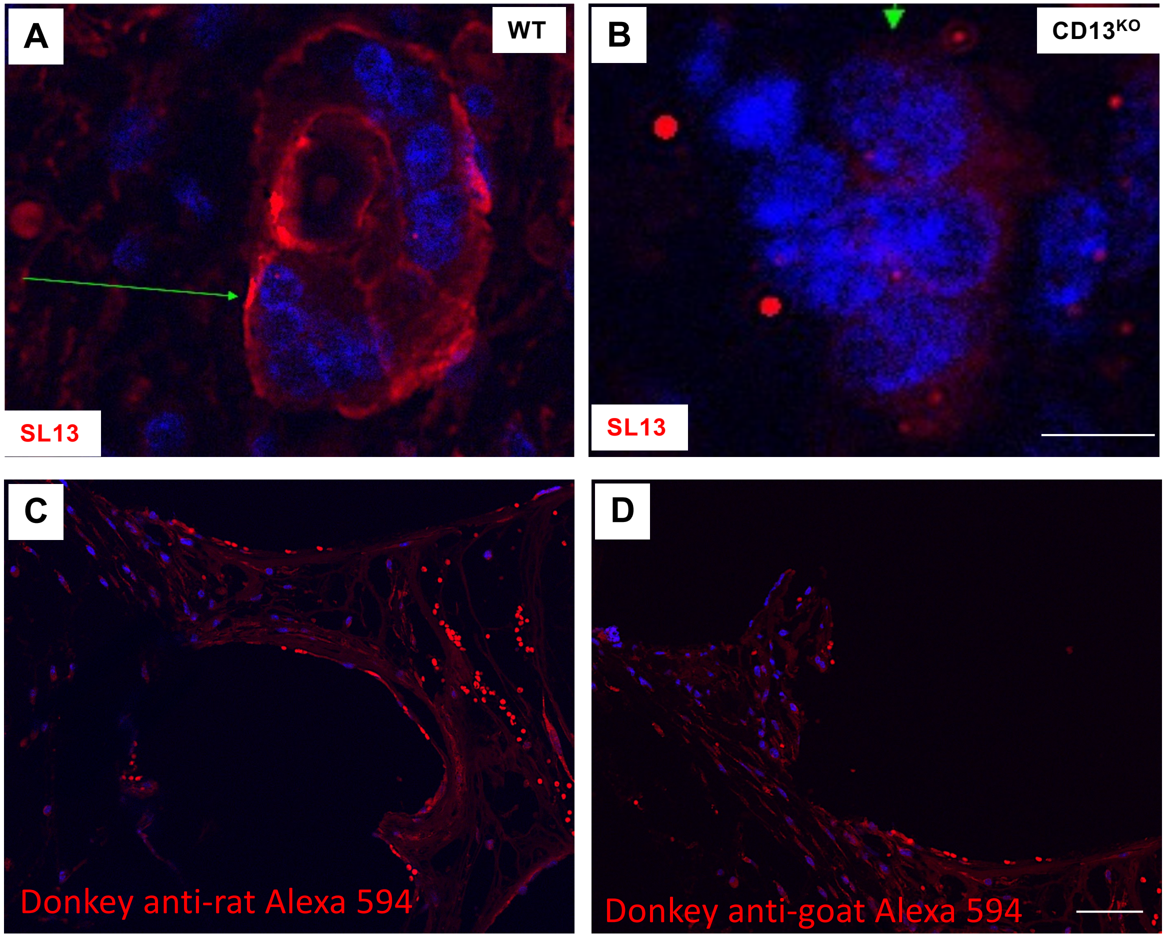

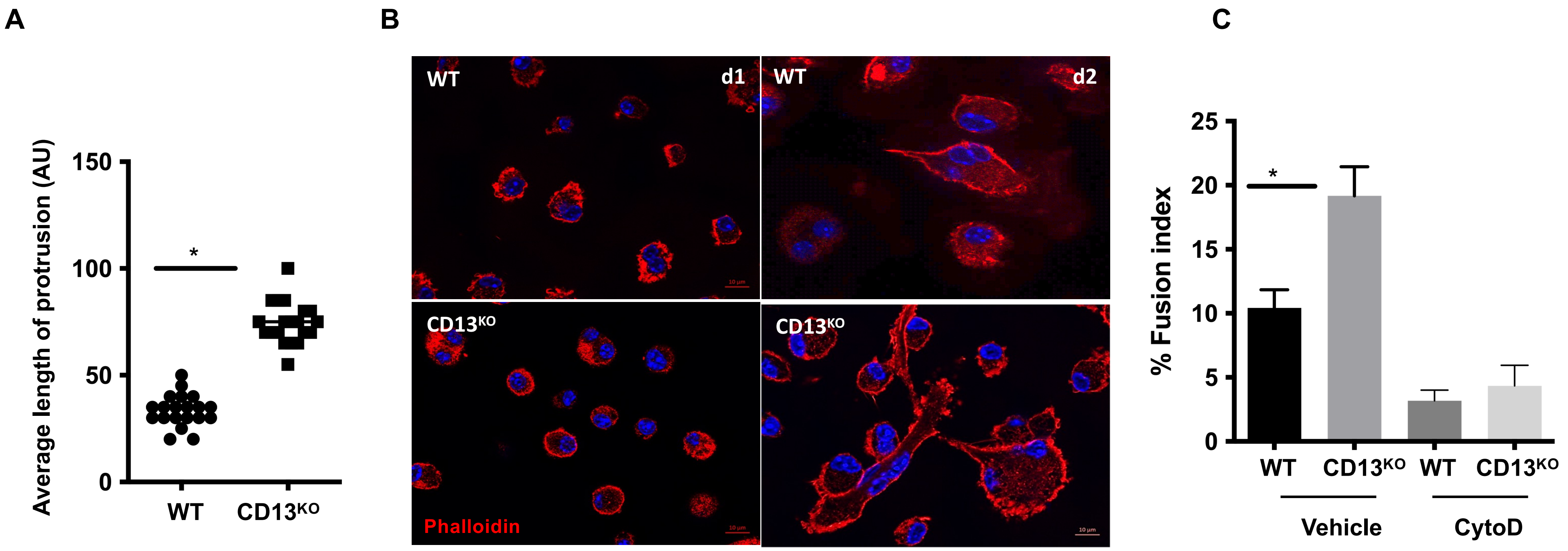

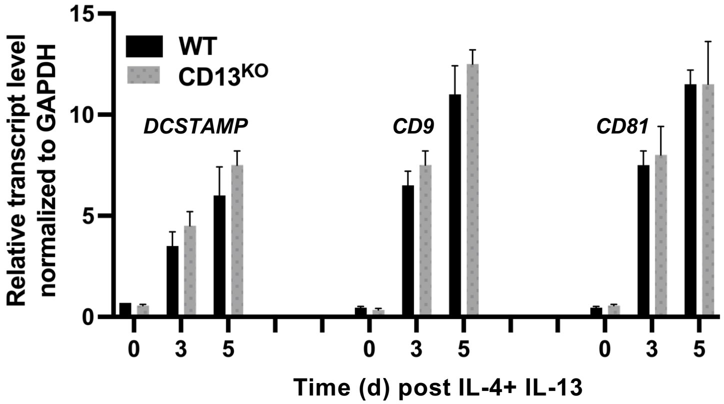

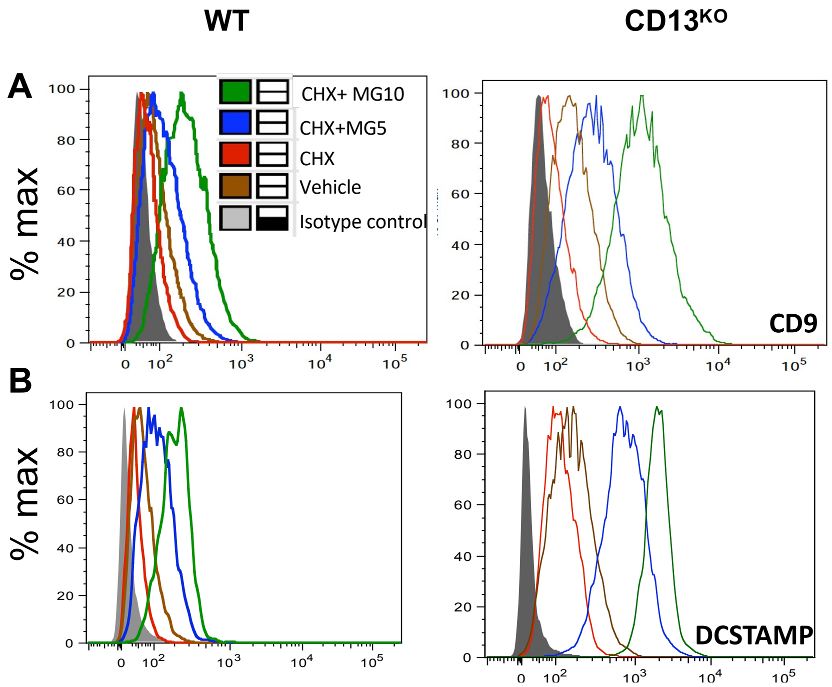

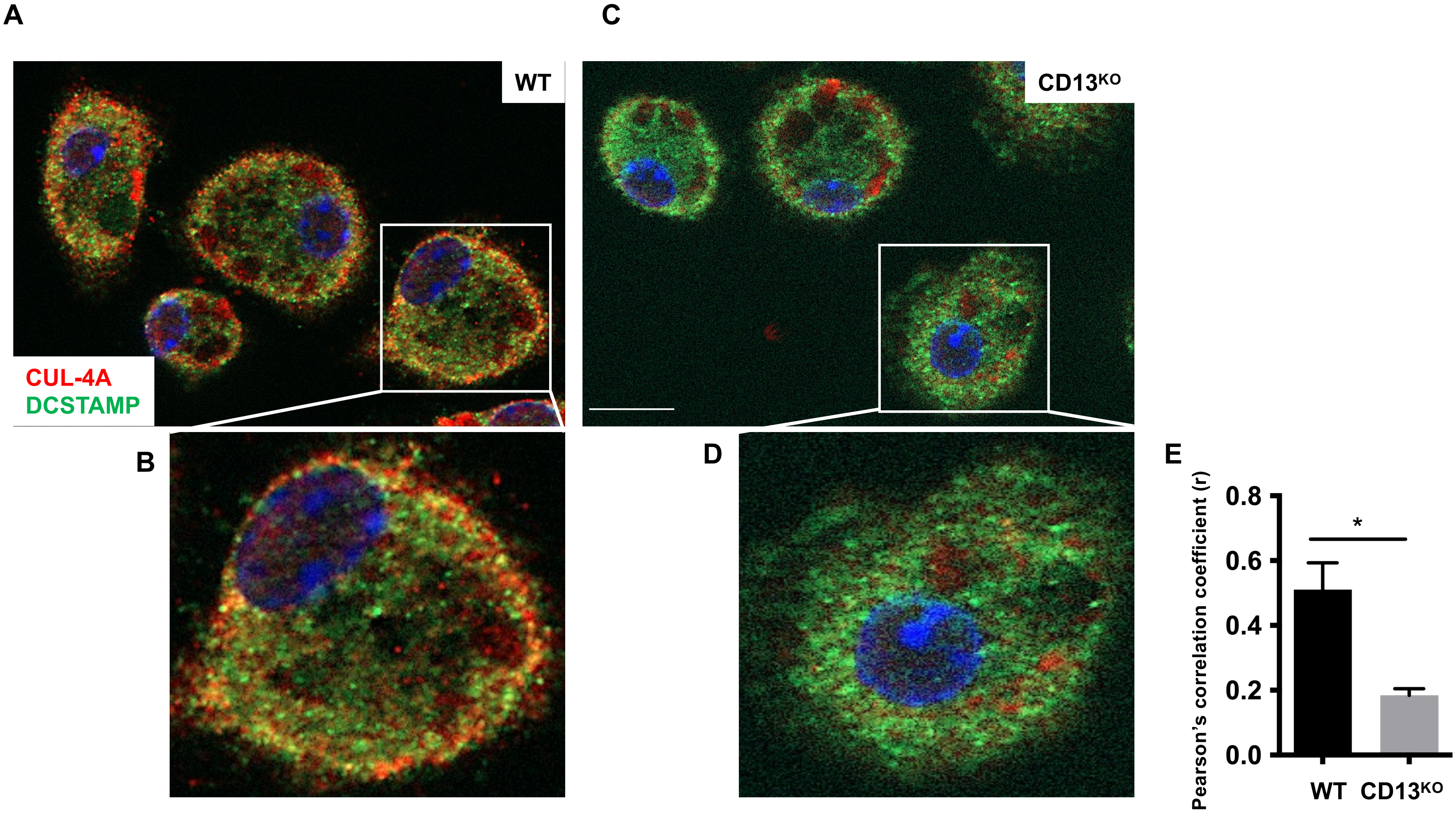

**FIG S7**

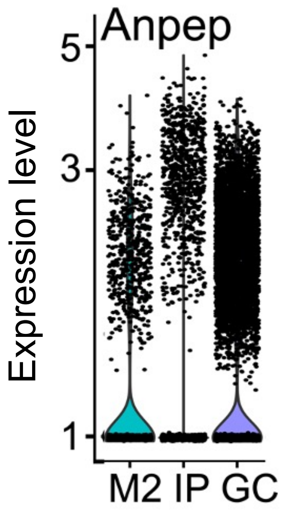

### Supplemental Figure 1

FIG S1

A

Live (CD3/B220/Nk1.1)⁻ CD11b lo/- CD115 hi Ly6G+ BM progenitor cells

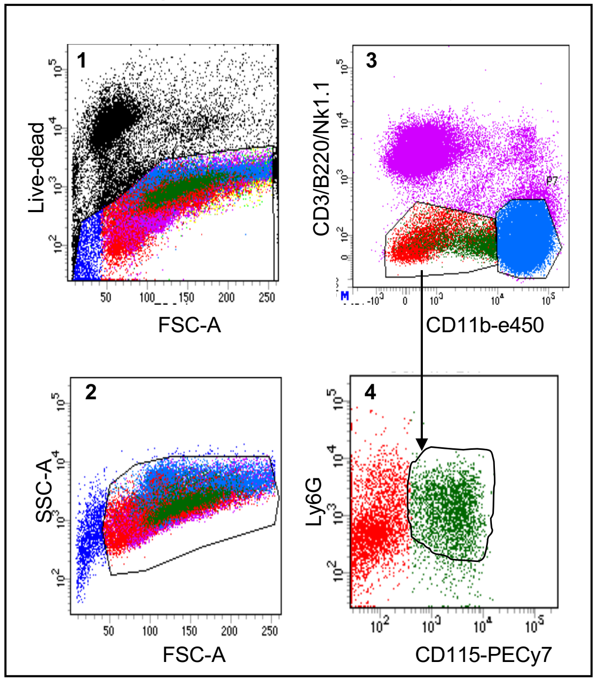


B

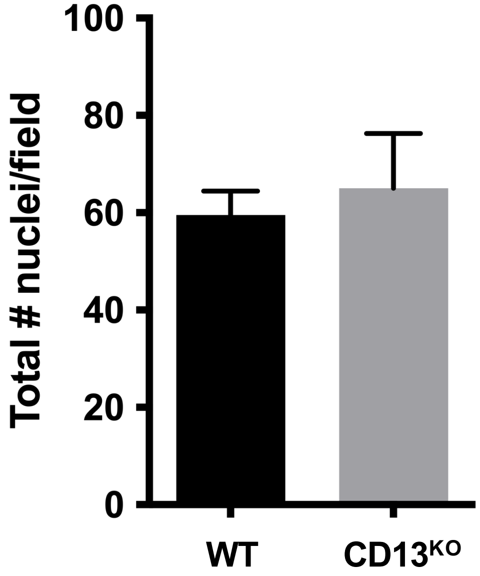

### Supplemental Figure 2

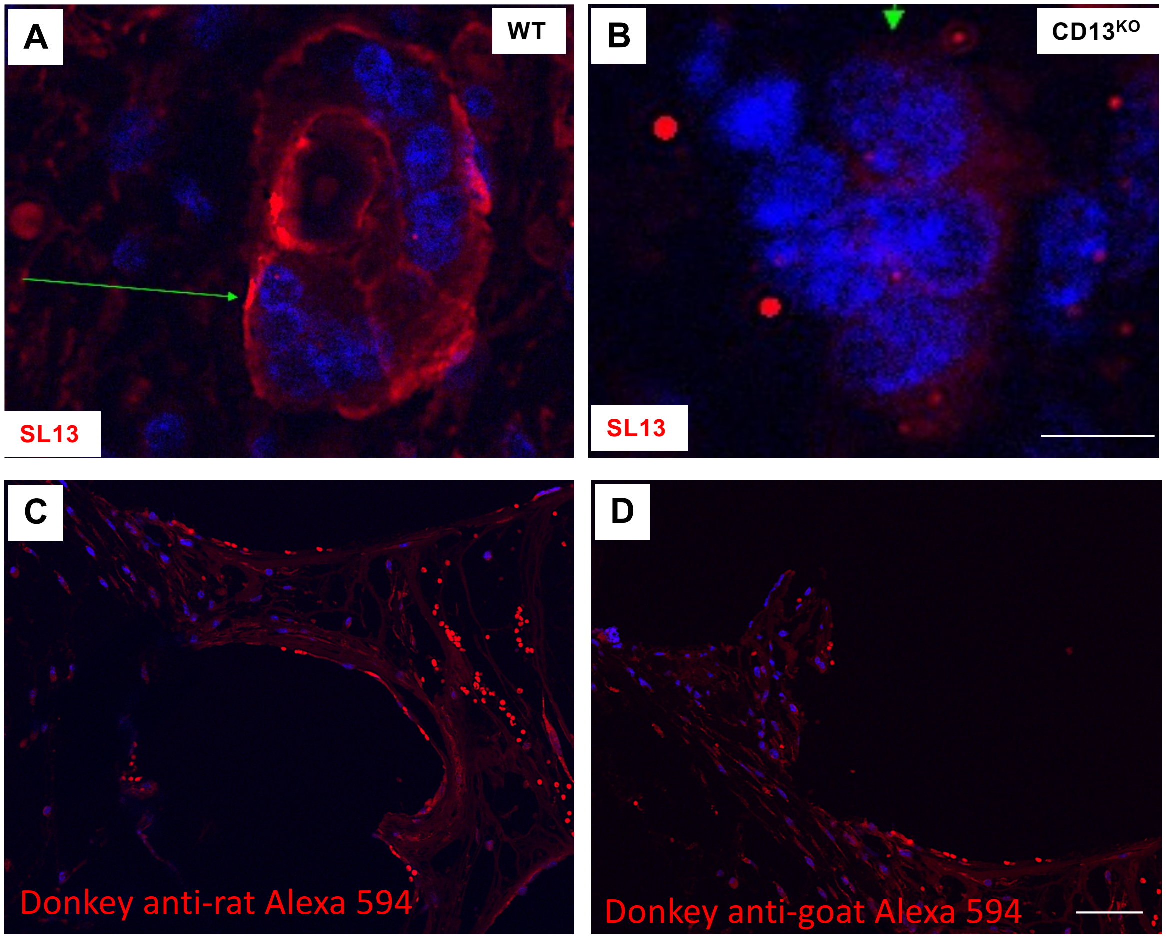

### Supplemental Figure 3

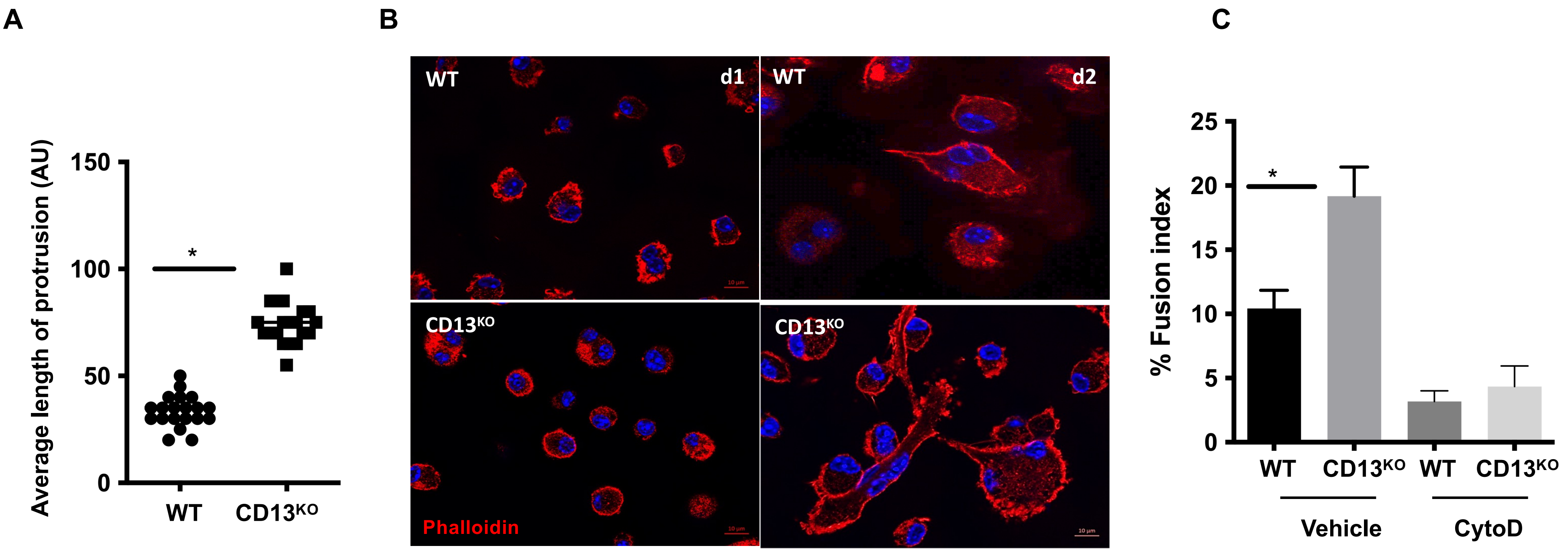

### Supplemental Figure 4

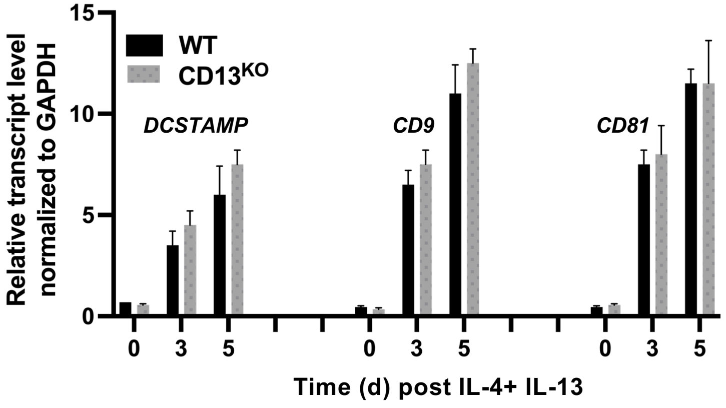

### Supplemental Figure 5

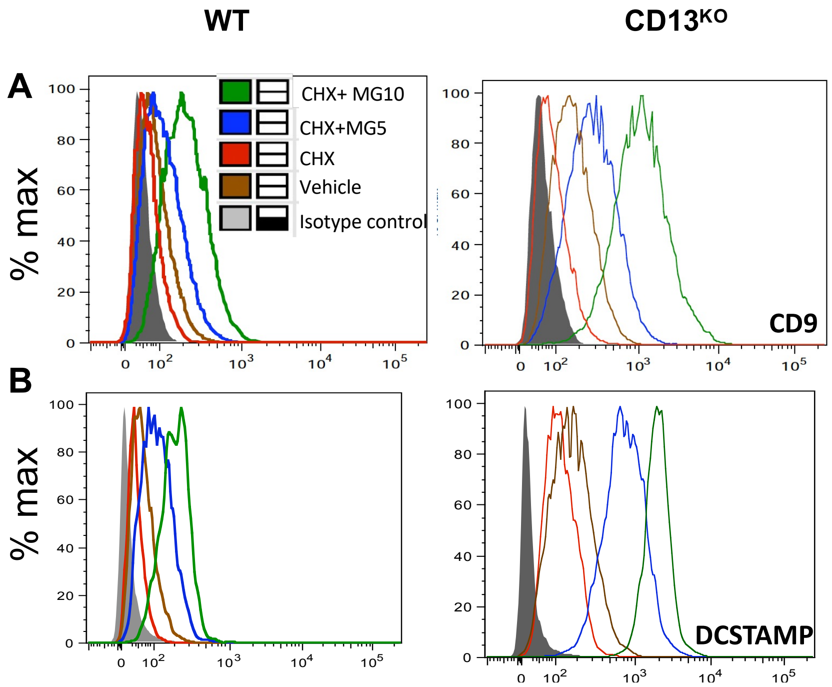

### Supplemental Figure 6

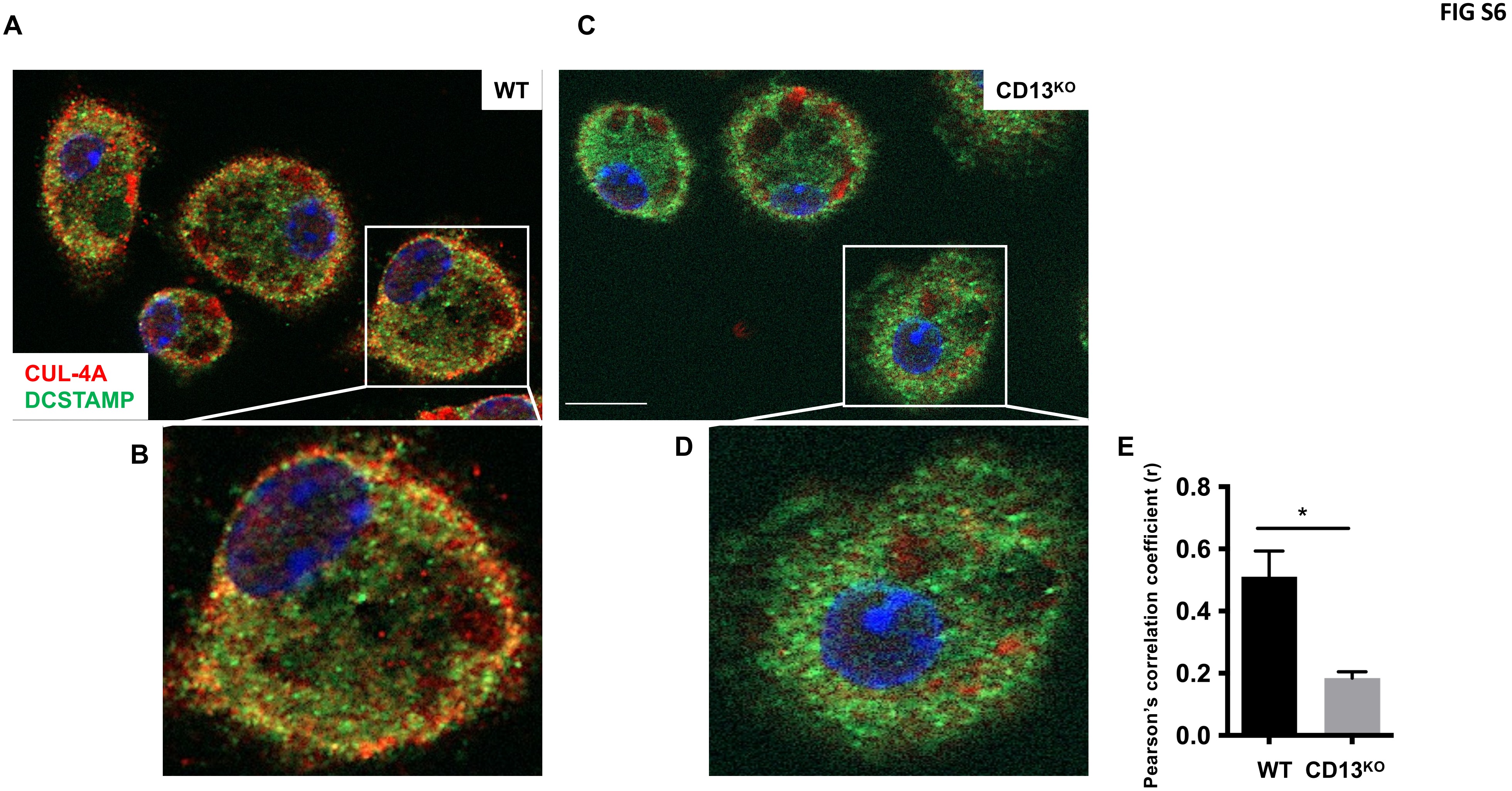

### Supplemental Figure 7

**FIG S7**

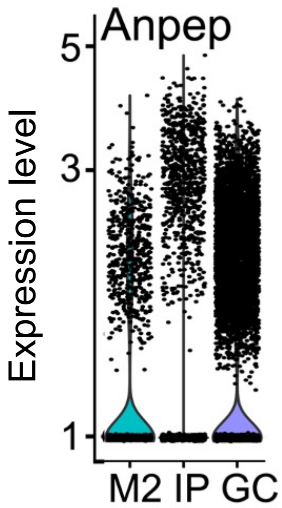
